# Sex Dimorphism in Expression of Immune Response Genes in Drosophila

**DOI:** 10.1101/2024.07.27.605461

**Authors:** MD Mursalin Khan, Rita M. Graze

## Abstract

Sex dimorphism in immunity is commonly observed in a wide variety of taxa and is thought to arise from fundamental life history differences between females and males. In *Drosophila melanogaster*, infection with different pathogens typically results in different modes of immune sex dimorphism, with male-or female-bias, likely due to specific disease-causing mechanisms of pathogens and host-pathogen interactions. Studies showed that some pathways, such as IMD and Toll, can explain these sex-dimorphic immune responses in Drosophila. However, it is unclear if sex differences in the immune response observed in *D. melanogaster* are conserved, even in closely related species. One window into identifying conserved and evolving sex differences in the immune response is to examine the sex-differential expression of immunity-related genes. Here, we aim to understand whether two closely related species, *D. melanogaster* and *D. simulans*, show conserved sex dimorphism in innate immunity, focusing on associated changes in gene expression in response to infection with a gram-negative bacterium, *Providencia rettgeri*. Survival, bacterial load, and bacterial load upon death (BLUD) were investigated to assess overall sex differences. *D. melanogaster* females and males differed significantly in survival, whereas *D. simulans* did not. Enrichment analyses revealed that both sexes and species upregulate genes involved in similar immune-related biological processes, but downregulated groups differed. We identified conserved sex differential gene expression of genes in the bacterial infection response pathways IMD, Toll, Jak/STAT, their regulators, and other immune-related gene classes (e.g., BOMs), as well as sex and species differences. In *D. melanogaster,* the effector antimicrobial peptides (AMPs) regulated by IMD were more highly upregulated relative to *D. simulans* in both sexes. Moreover, *D. melanogaster* females uniquely initiated high levels of gene expression that were involved in negative feedback mechanisms that controlled the overstimulation of IMD. Genes in the Toll pathway were also sex-differentially expressed with a higher level of upregulation in *D. melanogaster*. Remarkably, comparing expression across species, we find that *D. simulans* likely employs both the conventional peptidoglycan recognition-driven PRR-SPE-Spz pathway and the microbial protease recognition-based Psh-dependent activation of Toll; in contrast, *D. melanogaster* appears to solely rely on the PRR-SPE-Spz pathway in this context. In summary, our findings indicate that sex differences are conserved in both species for the majority of upregulated genes, while downregulation patterns and specific gene subsets show notable differences between sexes or species.

## 1. Introduction

Sex dimorphism influences a diverse set of biological processes, affecting nearly all aspects of an organism’s biology (Morrow 2015; Millington and Rideout 2018; Lopes-Ramos *et al*. 2020). Sex dimorphism of easily observable characteristics such as external coloration or anatomy has been widely described and studied, but dimorphism also extends to less readily apparent traits, including immune responses, infection outcomes, and disease prognosis (Carton et al. 1992; Fischer et al. 2015; Morrow 2015; McDonald et al. 2021; Wilkinson et al. 2022). Sex-biased immune responses have been observed at different stages of life history and may also be present in local tissue-level responses such as in the brain, intestinal layer, and hemolymph (Mackenzie et al. 2011; Deng and Jasper 2016; Millington and Rideout 2018; Belmonte et al. 2020; Khodursky et al. 2020; Wilkinson et al. 2022). The magnitude of sex differences may be mediated by the involvement of distinct immune compartments, such as hemocytes, melanization, epithelial immunity, and systemic immunity, depending on the pathogen and route of infection (Belmonte et al. 2020; Yu et al. 2022). At a mechanistic level, this may be explained by regulatory differences in key pathways, such as the Toll and IMD pathways, or more broadly by sex dimorphism of key tissues (Duneau et al. 2017b; Vincent and Dionne 2021). Even though immunological differences between sexes are observed frequently and impact fundamental mechanisms underlying immunity, these differences have received relatively little attention, including in model organisms and human studies (Morrow 2015; Klein and Flanagan 2016; Belmonte et al. 2020; Wilkinson et al. 2022).

Sex is a central axis of variation, explaining a significant amount of variation within species and differences between them (Bateman, 1948; Darwin, 1871; Lande, 1980). Sex differences have been observed in the intensity of the response against pathogens, specific outcomes of infections, and susceptibility, resistance, and tolerance to pathogens (Belmonte et al., 2020; Fischer et al., 2015; Klein & Flanagan, 2016). The evolution of differences in the immune response between males and females is most frequently attributed to differences in reproductive strategies and life history traits (Lande 1980; Zuk 2009; Morrow 2015). For instance, females typically invest more in offspring production and care, leading to the prediction that more robust immune responses will be associated with higher fitness in females (Lande 1980; Zuk 2009; Morrow 2015). Conversely, males may invest more in traits that enhance competitiveness, potentially at the expense of immune function (Lande 1980; Zuk 2009; Morrow 2015). Behavioral differences between males and females may also impact the frequency, intensity, and timing of exposure to various challenges such as viral, bacterial, fungal, and chemical, which could contribute to the evolution of sex differences or environmentally induced sex differences in immunity observed in natural contexts (Oertelt-Prigione 2012; Asahina 2018). In addition, other processes may intersect with sex dimorphism in immunity, contributing to variation within populations. For example, emerging work has detected sex differences in the aging of immune tissues, immune senescence, and inflammation (Mondal and Rai 1999; Metcalf et al. 2020; Abancens et al. 2020; Shepherd et al. 2021). Differences in food intake and diet are also observed between males and females and could modulate immunity in a sex-differential fashion (Oertelt-Prigione 2012; Asahina 2018; Rai et al. 2023).

Hormone signaling pathways, including sex hormones, have a role in the transcriptional regulation of the immune genes (Graze et al. 2018; Gal-Oz et al. 2019; Shepherd et al. 2021). Moreover, previous studies found that steroid hormones can directly regulate the function of the immune cells, such as the absence of ecdysone led to the inability of the macrophage-like cells to phagocytize bacteria in Drosophila (Regan *et al*. 2013). The peptide hormone IIS/TOR signaling disruption also reported the sex-differential immune gene expression in Drosophila (Graze *et al*. 2018). However, hormone signaling likely does not fully explain sex dimorphism in immune responses, as sex differences in immune response have been observed even at sexually immature life stages (Klein and Flanagan 2016; Shepherd et al. 2021). Sex chromosomes and dosage compensation mechanisms can also contribute to the differential expression of immune genes as well as the function of immune cells (Scofield et al. 2008; Libert et al. 2010). For instance, in mammals, the distinct composition of sex chromosomes can lead to female-specific mosaicism due to cell-specific X chromosome inactivation and may influence the expression of immune genes (Klein and Flanagan 2016; Pinheiro and Heard 2017). In humans, significant proportions of primary B lymphocytes, monocytes, and plasmacytoid dendritic cells express TLR7 on both X chromosomes by escaping X inactivation of XX individuals (Libert et al. 2010; Souyris et al. 2018; Heintze 2018). Overall, multiple aspects of sex determination and development impact sex dimorphism in immunity and often interact in complex ways.

Drosophila is a crucial model in the study of development and physiology and is growing as a model for sex differences in body size, aging, stress responses, and disease progression (Rideout et al. 2015; Millington and Rideout 2018; Belmonte et al. 2020). As a model organism for innate immunity, Drosophila showcases sex differences in both the cellular and humoral responses (Reviewed in (Belmonte *et al*. 2020)). These two central components contribute to sex-specific immune responses depending on the type of pathogen and the context of the infection (Hoffmann 2003; Lazzaro and Little 2009; Millington and Rideout 2018; Belmonte *et al*. 2020). Thus, *D. melanogaster* exhibits sexual dimorphism in the outcome of infectious diseases (Vincent and Dionne 2021; Rai *et al*. 2023; Khan and Graze 2024).

The Drosophila immune system comprises various cellular and humoral components that work together to recognize, signal proliferation of, and eliminate pathogens (Hoffmann 2003; Vlisidou and Wood 2015; Yu *et al*. 2022). One of the key cellular components of the *Drosophila* innate immune system is a specialized cell type called the plasmatocyte, the functional equivalent of mammalian macrophages/monocytes, which accounts for around 95% of the circulating hemocytes (Evans *et al*. 2003; Vlisidou and Wood 2015; Banerjee *et al*. 2019). Plasmatocytes can engulf and destroy invading microorganisms and maintain communication with the humoral response, such as response to antimicrobial peptides (AMPs) binding (Evans *et al*. 2003; Vlisidou and Wood 2015; Banerjee *et al*. 2019). The rest of the hemocytes (5%) primarily consist of crystal cells that produce phenol oxidase (PO) for melanization, primarily as an immune response involved in wound healing (Evans *et al*. 2003; Dudzic *et al*. 2015). Finally, upon pathogenic attack, fruit flies generate lamellocytes, which are larger than plasmatocytes and specialize in the encapsulation of large pathogens such as wasp eggs (Evans *et al*. 2003; Vlisidou and Wood 2015).

In the humoral response, the most critical aspect of the *Drosophila* innate immune response is the consistent production of small peptides called antimicrobial peptides (AMPs) and their assistance in recruiting additional immune cells such as plasmatocytes (Lai and Gallo 2009; Vlisidou and Wood 2015; Yu *et al*. 2022). Upon immune signal activation, AMPs are produced in the fat body and released into the hemolymph (Tzou *et al*. 2000; Hoffmann 2003). Due to their positive charge, AMPs are attracted to microbes with negatively charged cell membranes, where they embed in the hydrophobic region and cause membrane destabilization, resulting in cell death (Lai and Gallo 2009; Lin *et al*. 2020). Moreover, certain tissues, such as the trachea, midgut, oviduct, spermatheca, ganglia, and a subpopulation of hemocytes, produce AMPs in response to local infection (Tzou *et al*. 2000; Lai and Gallo 2009). In general, the level of AMP expression directly reflects the strength of the immune response (Lai and Gallo 2009; Lin *et al*. 2020).

The core response is initiated by pattern recognition receptors (PRRs) that recognize PAMPs, for example, specific molecular patterns on the surface of microorganisms, and initiate a signaling cascade that leads to the production of AMPs via the IMD and Toll pathways (Tanji and Ip 2005). The IMD and Toll pathways are exceptionally well studied in this system (Valanne *et al*. 2011; Myllymäki *et al*. 2014). They are homologous to the vertebrate TNF and Toll-like pathways, respectively (Tanji and Ip 2005; Yu *et al*. 2022; Valanne *et al*. 2022). In general, the IMD pathway is activated upon the detection of peptidoglycan produced by gram-negative bacteria, whereas the Toll pathway responds to the peptidoglycan of most gram-positive bacteria, and fungal cell wall components such as β-(1,3)-glucan (Tanji and Ip 2005; Valanne *et al*. 2022). Moreover, the Drosophila STING pathway activates upon detection of bacterial cyclic dinucleotides (CDNs) to initiate an innate immune response by triggering the NF-κB transcription factor Relish of IMD pathway to produce AMPs (Martin *et al*. 2018). Of note, a second cascade besides PRR-based activation, the danger signal cascade, can activate the Toll pathway (Ligoxygakis *et al*. 2002; Gottar *et al*. 2006; Chamy *et al*. 2008). The danger cascade initiates upon the detection of the abnormal activity of bacterial or fungal proteases (danger signal) and activates the Toll pathway by using Clip-Serine Protease Persephone (Psh) via Psh-SPE-Spz or Psh-Protease-Spz (SPE independent) cascades (Gottar *et al*. 2006; Yamamoto-Hino and Goto 2016; Issa *et al*. 2018; Dudzic *et al*. 2019). Upon activation, each pathway launches a rapid and efficient transcriptional response to infection via nuclear translocation of NF-κB family transcription factors, Dif and Dorsal (Tanji and Ip 2005; Valanne *et al*. 2022).

Recent studies have reported that *Drosophila* males and females differ in response to localized and systemic bacterial, fungal, viral, and parasitic infections (Reviewed in (Klein and Flanagan 2016; Belmonte *et al*. 2020)). In many cases, for example, Kalithea (viral), *Candida albicans* (fungal), *Tubulinosema ratisbonensis* (microsporidial), and *Serratia marcescens* (bacterial), there was a bias toward females in the effectiveness of the immune response as measured by survival (Kutch and Fedorka 2017; Palmer *et al*. 2018; Franchet *et al*. 2019; Chowdhury *et al*. 2019). At the same time, this pattern is not consistently observed even among similar types of pathogens; male flies infected with *Beauveria bassiana* (fungal), *Providencia alcalifaciens* (bacterial), and *Enterococcus faecalis* (bacterial) have higher survival rates relative to females (Taylor and Kimbrell 2007; Le Bourg 2011; Duneau *et al*. 2017b; Shahrestani *et al*. 2018). Collectively, these studies, along with others, have also revealed considerable context dependency. Sex dimorphism in immunity can be influenced by various factors such as hormonal regulation, environmental conditions, dietary habits, and metabolic processes (Leech *et al*. 2019; Shepherd *et al*. 2021; Rai *et al*. 2023). Conditional impacts are especially evident when the same pathogen shows different directions of sex dimorphism under different experimental conditions, either in different laboratories or in the same study with different treatments (Rai *et al*. 2023). For example, two independent studies of *Pseudomonas aeruginosa* infections demonstrate opposite-sex bias (Vincent and Sharp 2014; Chowdhury *et al*. 2019). Moreover, recent findings suggest that sex-determination pathway genes such as *transformer (tra)* might influence immune sex dimorphism in Drosophila (Khan and Graze 2024).

Several recent reviews have highlighted a significant gap in our understanding of sex differences in immunity, particularly in adult animals and in the context of evolutionary biology (Beery and Zucker 2011; Belmonte *et al*. 2020). Further, there is a relative lack of comparative studies to understand which aspects of sex dimorphism in the immune response are conserved. Comparative studies are also crucial for understanding the evolutionary forces influencing sex differences across the species. For instance, comparative studies of closely related species can help to expound whether observed sex differences are lineage-specific and/or adaptive (Hill-Burns and Clark 2010; Khodursky *et al*. 2020; Nanni *et al*. 2023).

In *D. melanogaster*, both the IMD and Toll pathways play pivotal roles in immune dimorphism (Duneau *et al*. 2017b; Vincent and Dionne 2021). The Toll pathway and its regulators may be responsible for a significant portion of immune sex dimorphism in *D. melanogaster* (Duneau *et al*. 2017b). In addition, regulation of the IMD pathway also plays a role in sex differences (Vincent and Dionne 2021). While this is a growing area of study, there are many remaining questions regarding the complex molecular basis of sex dimorphism in immune responses and the degree to which the role of the Toll and IMD pathways in sex dimorphism is conserved. The intricate crosstalk among evolutionarily conserved immune pathways, including Toll, IMD, Jak/STAT, and JNK, introduces further complexities to the investigation of sex dimorphism (Dudzic *et al*. 2019; Tafesh-Edwards and Eleftherianos 2020; Alejandro *et al*. 2022). Overall, a comprehensive understanding of sexual dimorphism in immunity can provide valuable insights into the complex interplay among evolution, diversity, immunity, and sex, which also has important implications for the treatment and prevention of disease.

To comprehensively characterize sex dimorphism in immunity in *Drosophila*, we conducted a comparative study, profiling outcomes of infections and gene expression in unmated-adult females and males in two species, *D. melanogaster* and *D. simulans*. This study allows a direct comparison of the response to infection in *D. melanogaster*, where most of the immunological studies have been completed, and *D. simulans*, a relatively well-studied and closely related species of the same sub-group (Figure 1A) (Bhutkar *et al*. 2007; Paris *et al*. 2013). In addition, to the best of our knowledge, this is the first time bacterial infection in *D. simulans* has been reported for both sexes, and it is the first comparative immune response study with *D. melanogaster.* This work identifies sex differences that are in common, as well as divergence of sex differences in immune gene expression, laying the groundwork for an improved understanding of the evolutionary context of sex dimorphism in the immune response.

**Figure 1:**
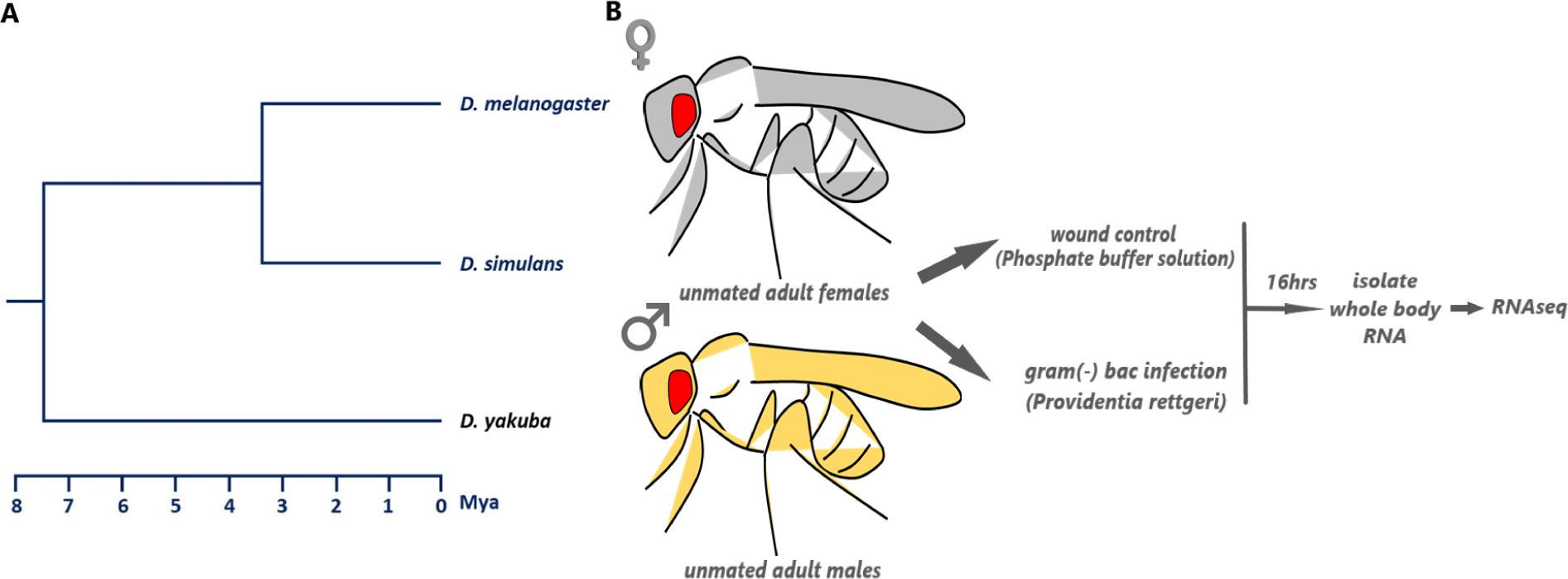
Phylogenetic relationship and experimental design. **A.** Phylogeny (represented as a schematic diagram) of *D. melanogaster* and *D. simulans*, based on (Bhutkar et al. 2007; Paris et al. 2013; Li et al. 2022). *D. melanogaster* and *D. simulans* were used for the experiments, and *D. yakuba* is shown as an outgroup. Mya: millions of years ago **B.** Experimental Design. The unmated adult flies were infected with *P. rettgeri* (PRET) for the treatments and phosphate buffer solutions (PBS) for wound control. RNAseq was performed from RNA isolated from pooled flies (whole body) at 16hpi in four replicates.

## 2. Methods and Materials

### 2.1 Fruit fly stocks and sample collection

*D. melanogaster* and *D. simulans* were reared on a modified version of the Bloomington Drosophila Stock Center (BDSC) standard cornmeal medium recipe (33 L H_2_O, 237 g agar, 825 g dried deactivated yeast, 1560 g cornmeal, 3300 g dextrose, 52.5 g Tegosept in 270 mL 95% ethanol and 60 mL propionic acid). Flies were reared at 25°C with a 12:12 light: dark cycle. Density-controlled crosses were set up for each species, *D. melanogaster* (Canton S, Gift Michelle Arbeitman) and *D. simulans* (Dsim w[501], 14021-0251.195, National Drosophila Species Stock Center, Cornell University); newly emerged progeny was collected as virgins and separated by sex. Female and male individuals were isolated in standard vials, and all experimental flies were unmated, adult, and aged 5-9 days prior to infection.

### 2.2 Experimental treatment

A strain of the gram-negative *Providencia rettgeri* species isolated from wild-caught *D. melanogaster* (*P. rettgeri* strain: Dmel, Gift Brian Lazzaro) was used as an infectious agent, which is known as moderately virulent (Juneja and Lazzaro 2009; Galac and Lazzaro 2011; Duneau *et al*. 2017b). In this study, stationary phase *P. rettgeri* bacterial culture (OD600nm = 1.2) was used to inject bacteria into the thoracic region (soft spot near the wings) of the fly to introduce around 1200-1500 cells directly into the hemolymph using the needle prick method (Figure 1B; following (Khalil *et al*. 2015)). For control flies, sterile phosphate buffer solution (1x PBS) was injected as a wound-only control (Figure 1B).

### 2.3 Bacterial culture preparation

Glycerol stocks of *P. rettgeri* were stored at −80 °C prior to use. Stocks were used to streak LB-Agar plates (Teknova, California) and incubated at 37°C for 12 hours to isolate single colonies. A single colony was then transferred to 5 mL Lysogeny broth LB-Broth (Teknova, California) and incubated at 37°C at 220 rotation per minute (rpm) for 9-10 hours. Stationary phase bacteria were centrifuged at 4000 rpm for 3 minutes to collect bacterial pellets. Pellets were washed with sterile 1x PBS using pipette aspiration (5 times) to remove residual LB Broth. After five washes, the pellet was resuspended with sterile 1x PBS to achieve OD600 nm of 1.2.

### 2.4 Bacterial injection of flies

The flies were anesthetized by CO_2_ during the injection. The injections were made in the thoracic region beside the wings in the soft spot using the needle pricking techniques previously described by Khalil et al. 2015 to introduce approximately 1200-1500 viable bacteria into each fly. This approach introduces bacteria directly into the hemolymph. The fly vials were kept horizontal to allow recovery until the flies were completely active.

### 2.5 Survival rate

For each sample type, the survival assay was performed in 5 replicates, each of which contained 30 flies. Survival was scored at 2, 6, 12, 18, 24, and 36 hours after bacterial injection. Flies were scored as dead if the fly was on its back and was not moving or could not turn by itself (Duneau *et al*. 2017a). Scoring ceased when all the flies were dead in any of the experimental groups or after the 36-hour time point. Survival analysis was conducted in R using the package “survival” (Therneau 2023). Kaplan-Meier survival curves were generated to visualize the survival proportion. The Cox regression was performed to determine the statistical difference among experimental groups (Therneau and Grambsch 2000).

### 2.6 Bacterial load time series

Bacterial load was assessed for each sample in six replicates, with each replicate consisting of one fly at each time point. The bacterial load was determined at 4, 8, 12, 16, 20, and 24 hours after injections, with six replicate measurements following Duneau et al. (Duneau *et al*. 2017a). A single fly per replicate measurement was homogenized in 200 µL 1X PBS using a sterile pestle. The homogenized sample (10 µL) was transferred to a 96-well plate and diluted with 90 µL 1X PBS. From the first dilution (1/10), 10 µL of the solution was transferred successively, and serial dilution was carried out 8 times. Subsequently, 20 µL of the dilution 1:10^2,^ 1:10^4^, 1:10^6^, and 1:10^8^ were spread on LB agar plates to count CFU. Plates were incubated at 37 °C for 12-14 hours to count the CFU before gut microbes typically form colonies that occur at approximately 36 hours (Khalil *et al*. 2015). To test the differences between infected and control animals for each time point and to get the differences across groups, the linear model was used (Sparks 2017).

### 2.7 Bacterial load upon death (BLUD)

The bacterial load upon death was determined for each sample in four replicates, with each replicate comprising three flies per pool. For the BLUD experiment, flies were determined to be dead flies if the fly was on its side or back without any movement (Duneau *et al*. 2017a). A single fly was homogenized in 200 µl of 1x PBS within 30 minutes of determining death using a sterile pestle. Serial dilutions and CFU counts were obtained, as described in the previous section. To find the differences in groups for each time point, the linear model (CFU ∼ Groups) was fit and pairwise differences tested for species and sex comparisons.

### 2.8 RNA extraction and prep

Four replicate pools of 5–9 day old flies with 20 flies per pool were collected for each sex and each group. Only live flies were used for RNA isolation. Replicate pools of wound control and bacteria-infected flies were flash-frozen in liquid nitrogen 16 hrs post-injection (hpi). The frozen samples were stored at −80°C prior to total RNA isolation. The 20 flies (per pooled sample) were homogenized entirely in 700 µL of TRI-Reagent using a pellet pestle motor and RNase-Free pellet pestles in a 2 mL microcentrifuge tube. Total RNA was extracted using the Direct-zol RNA Miniprep Plus Kit (Zymo Research) according to the manufacturer’s instructions. The TURBO DNase-free kit (Thermo Fisher Scientific) was used to remove genomic DNA contamination following the manufacturer’s instructions. The concentration was determined using Nanodrop 2.0 (Thermo Fisher Scientific), and RNA integrity Number (RIN) was measured by using a TapeStation system with an RNA ScreenTape Analysis kit (Agilent Technologies).

### 2.9 TagSeq library prep

3’ TagSeq was performed at the University of Texas Genomic Sequencing and Analysis Facility, Austin (UT-GSAF), based on the protocols from Lohman et al. (2016) and Meyer et al. (2011) (Lohman et al., 2016; Meyer et al., 2011). Libraries were quantified using the Quant-it PicoGreen dsDNA assay (ThermoFisher) and pooled equally for subsequent size selection at 350-550 bp on a 2% gel using the Blue Pippin (Sage Science). The final pools were checked for size and quality with the Bioanalyzer High Sensitivity DNA Kit (Agilent Technologies), and their concentrations were measured using the KAPA SYBR Fast qPCR kit (Roche). Libraries were sequenced on a NovaSeq 6000 (Illumina), 100 bp single-end reads, with 5-10 million reads per replicate (File S1).

### 2.10 Preprocessing and alignment

Single-end 100 bp reads were generated using 3′ Tag RNA Seq for each experimental group with 4 replicates each. The total raw reads generated by 3′ Tag RNA Seq was between 5.7 and 8.4 million (M) reads per sample, with an average of 6.99 M reads for *D. melanogaster* and 7.16 M reads for *D. simulans* (File S1). The quality assessment of raw data was performed by FastQC (Babraham Institute, University of Cambridge) (Anders 2010). The FastQC analysis showed that all 32 samples had high quality throughout, and no trimming was performed except the removal of the adapter by the sequencing facility (Trimmomatic; Bolger et al., 2014).

The reads were aligned against the respective *Drosophila* genomic references (FlyBase Releases: dmel_r6.47 for *D. melanogaster* and dsim_r2.02 for *D. simulans*) using the read alignment tool STAR (Gelbart *et al*. 1996; Gramates *et al*. 2022). The alignments were used to generate raw read counts for each gene in the reference genome annotation by using the “genomeDir”, “readFilesIn”, and “outSAMtype” options in STAR (Dobin *et al*. 2013). Gene expression levels were then quantified using the “quantMode GeneCounts” option of STAR (Dobin *et al*. 2013).

The mean uniquely mapped reads percentage for *D. melanogaster* samples was 74.48%, and for *D. simulans* samples, it was 76.81%, which is typical for 7 M reads of 3′ Tag RNA Seq (File S1) (Ma *et al*. 2019). Upon mapping, raw read counts of 17,633 genes in *D. melanogaster* and 15,266 genes in *D. simulans* were generated by STAR. Of 17,633 genes of *D. melanogaster* and 15,266 genes of *D. simulans*, 13,588 and 13,010 genes passed the detection cutoff, respectively (File S2-S5). Each experimental sample passed the quality control analysis prior to normalization and DEG analysis.

An edgeR-limma-voom workflow in R, was used to normalize raw counts (Log2CPM) and fit a cell means model, with defined contrasts used to determine differential expression between experimental groups, focusing on sex and species differences in gene expression resulting from infection (Robinson *et al*. 2010; Ritchie *et al*. 2015). Differences were classified as up or downregulated relative to wound control. The interaction of treatment by sex was tested to identify significant differences between the sexes, within each species (Law *et al*. 2014; Ritchie *et al*. 2015). A significant treatment by sex interaction term means that the sexes differ in the way that the gene expression for a gene changes in response to the infection. Additional tests for interactions were included to investigate species differences. In between-species comparisons, significant interaction terms for a gene indicate differences between species in the changes in gene expression observed in comparisons for a single-sex (treatment by species) or in comparisons between sexes (treatment by sex by species).

### 2.11 Differential gene expression analysis

Gene expression differences were first evaluated for each species individually to allow analysis of all expressed annotated genes in a species. Genes with fewer than 2 read counts (summed across samples within a group) were excluded from the analysis. Expression was estimated as the Log2CPM normalized counts per gene for *D. melanogaster* (mel) or *D. simulans* (sim), females (F) or males (M), with infection by *P. rettgeri* (pret) or PBS-only injected controls (PBS). Voom variance modeling was implemented in limma, with a means model and comparisons between groups tested using contrasts (Law *et al*. 2014; Ritchie *et al*. 2015). In *D. melanogaster,* the comparisons tested were mel_f_pret = mel_f_pbs and mel_m_pret = mel_m_pbs, the effect of treatment for each sex, and the interaction of treatment by sex mel_f_pret - mel_f_pbs = mel_m_pret - mel_m_pbs. Likewise, for the *D. simulans*, the corresponding tests were conducted. For the enrichment, gene class heatmaps, and pathway analysis in *D. simulans,* the annotated *D. melanogaster* orthologs were used (dmel_orthologs_in_dsim_Flybase_2017_05; File S13). An FDR-adjusted *p*-value (*p*-adj) cutoff of 0.05 was used for all tests of differential gene expression.

For comparisons between species, unique ortholog pairs were identified and included in the analysis. If two putative orthologs were found for one gene, then only one representative was kept for the analysis, meaning a single representative pair for each set of ambiguous homologs, for which orthology has not been clearly defined, was included in the analysis. As mentioned above, raw counts per gene were normalized using edgeR-limma-voom. For the comparison between species, normalization also included TMM density normalization of Log2CPM transformed counts. The contrast used to test the interaction of treatment by species in females is dmel_f_pret - dmel_f_pbs = dsim_f_pret - dsim_f_pbs, and the effect of treatment by species in males is dmel_m_pret - dmel_m_pbs = dsim_m_pret - dsim_m_pbs (Law *et al*. 2014). Additionally, contrasts were used to test the three-way interaction of treatment by sex by species ((dmel_f_pret - dmel_f_pbs) - (dmel_m_pret - dmel_m_pbs)) = ((dsim_f_pret - dsim_f_pbs) - (dsim_m_pret - dsim_m_pbs)) (Law *et al*. 2014; Ritchie *et al*. 2015). An FDR-adjusted *P*-value (*p-*adj) cutoff of 0.05 was used for all tests conducted. Unless otherwise mentioned, all the reported log2FoldChanges (log2FC) and interactions are significant at p-adj < 0.05.

### 2.12 Enrichment and pathways

The “clusterProfiler” R program package was used for gene ontology (GO) based gene set enrichment analysis (Wu *et al*. 2021). GO-Biological Process terms with adjusted *p*-value (FDR) less than 0.05 were considered significantly enriched. GO-BP annotations were performed separately for each contrast. The enrichment of DEGs in the Kyoto Encyclopedia of Genes and Genomes (KEGG) pathways was also tested using the “clusterProfiler” package in the R (FDR < 0.05). Expression differences of DEGs for selected pathways were visualized using “Pathvisio” and the wiki pathways plugin. The wiki pathway WP3830 was used to generate the Toll-IMD-Jak/STAT gene list and pathways.

### 2.13 Functional groups

Flybase was used to identify the functional groups/class gene list using key terms for Bomanin (BOMs), Drosophila immune-induced molecules (DIMs), Turandot (Tots), Thioester-containing protein (Teps), Insulin, Antimicrobial peptides (AMPs), Jun N-terminal kinase (JNK)/TNF-alpha-Eiger Pathway and Regulator, Positive and Negative Regulators of Toll, Positive and Negative Regulators of immune deficiency (IMD), and Spätzle (Spz) Activating Pathway with Regulators (Gelbart *et al*. 1996; Gramates *et al*. 2022). The Jak/STAT and melanization gene list was created manually from earlier studies and review articles (Tang 2009; Myllymäki and Rämet 2014; Yu *et al*. 2022). Heat maps of functional groups/classes were generated by “ggplot2” in the R program.

## 3. Results

Sex dimorphism is commonly observed in immune responses but is variable based on the pathogen, host, and other environmental factors (Fischer *et al*. 2015; Klein and Flanagan 2016). There is clear evidence of sex differences in immunity in *Drosophila melanogaster*, both at the physiological and molecular level, with a few genes identified as sex-differentially induced or repressed in response to infection (Taylor and Kimbrell 2007; Duneau *et al*. 2017b; Shahrestani *et al*. 2018). However, the range and causes of variation in sex dimorphism in immune responses, how and why these differences have evolved, and the underlying molecular mechanisms are unknown. This study aimed to understand if sex differences in the immune response are conserved between closely related species of *Drosophila* (Figure 1A). The study characterized the sex differences between unmated males and females associated with the immune response against a gram-negative extracellular bacterium, *P. rettgeri*, in *D. melanogaster* and its close relative *D. simulans*, by combining survival assays with RNA-seq-based gene expression estimation (Figure 1B). DEGs were analyzed within species, for all annotated genes, and between species for orthologs.

### 3.1 Sex dimorphism in survival

Fruit flies initiate rapid and efficient innate immune responses in response to pathogens that are crucial for survival, which influence the eventual outcome of bacterial infections. As one metric of relative effectiveness of the immune response, survival assays were conducted for females and males of *D. melanogaster* and *D. simulans* upon *P. rettgeri* infection over 36 hours at 6 to 12-hour intervals in five replicates where each replicate contained 30 flies (Figure 2). The survival assay demonstrated significant sex dimorphism in *D. melanogaster* (*p*-value < 0.001), but no significant differences in survival between the sexes were identified in *D. simulans.* This demonstrates that there are species differences in sex dimorphism in response to bacterial infection. Primarily, the initial stages (0-12hpi) showed no significant difference in survival between sexes for either species (Figure 2A). Nonetheless, survival started to differ by sex around 12hpi when the infection was expected to be established. At 18hpi, there were significant differences between males and females, where females exhibited higher survival than males in *D. melanogaster*. This difference in survival rate was persistent between sexes from 18hpi to 36hpi. Overall, the Kaplan Meier survival curve and Cox-regression model for *D. melanogaster* showed significant differences between males and females for *P. rettgeri* infection (Figure 2A).

**Figure 2:**
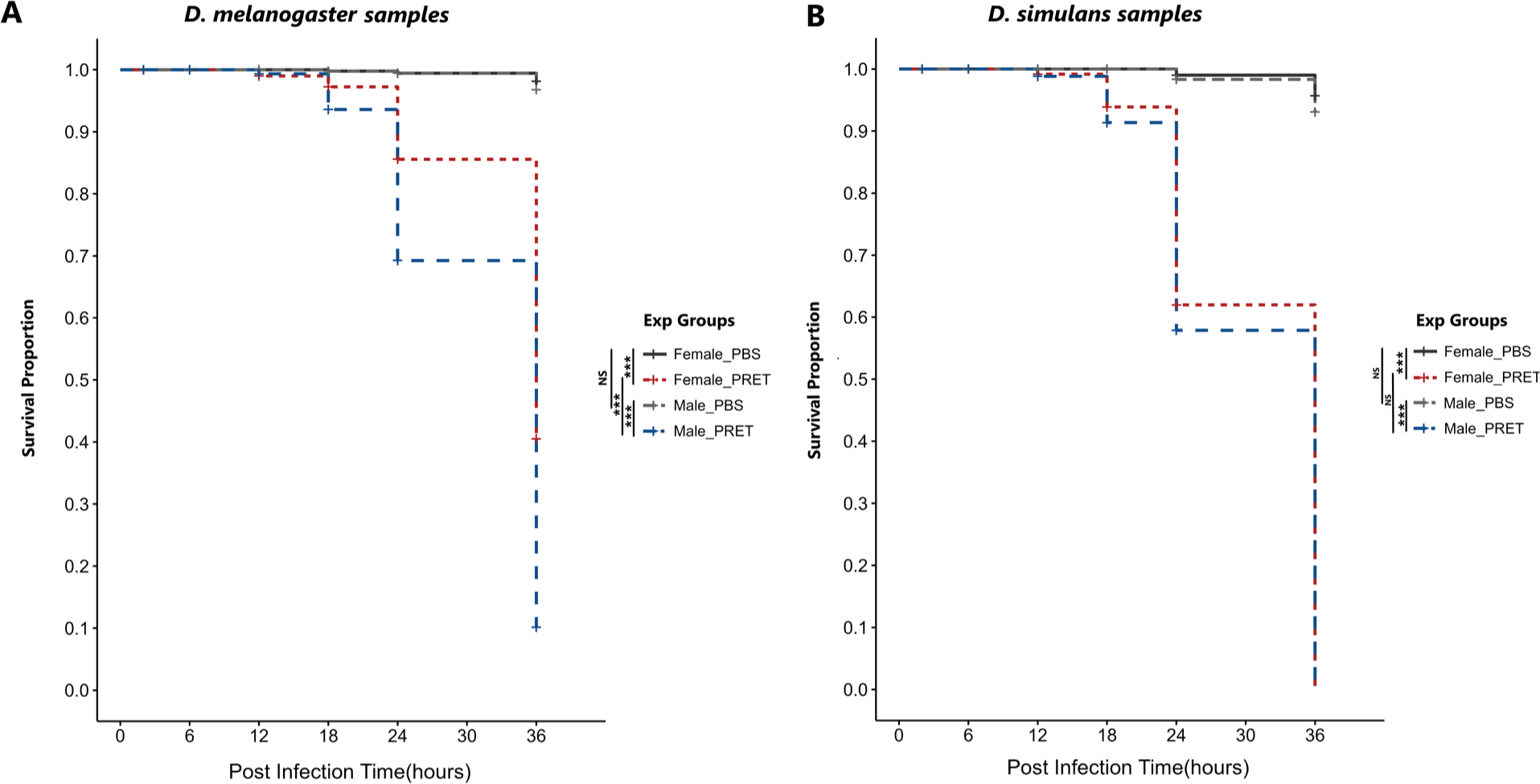
Determination of survival. **A.** Survival curve of *D. melanogaster* (Mel) upon infection with *P. rettgeri* (PRET) and wound control (PBS). **B.** Survival curve of *D. simulans* (Sim) upon infection with *P. rettgeri* (PRET) and wound control (PBS). The survival curve was generated with the Kaplan–Meier estimator, and the statistical analysis was performed using Cox regression. The asterisks (***) demonstrated the *p-*values < 0.001, and NS represented no significance. All the experimental flies of the *D. simulans* were identified dead at 36hpi for females and males.

For *D. simulans*, a sister species and member of the *D. melanogaster* subgroup, comparable results were observed in the initial stages of the infection (up to 12hpi) (Figure 2B). However, survival data began to diverge at 18hpi relative to *D. melanogaster.* In *D. simulans,* females showed no survival differences compared to males. While males and females continued to have no survival differences for 18hpi to 36hpi, the *D. simulans* sexes did not differ significantly in overall survival. Unlike *D. melanogaster* flies, all the *D. simulans* infected flies had died at 36hpi for both sexes (Figure 2B).

Comparing between species, female *D. melanogaster* had significantly higher survival after the establishment of the infection relative to *D. simulans* (*p*-value < 0.001). In all data points after infection initiation, *D. melanogaster* females consistently showed significantly higher survival rates than *D. simulans* females. Males of both species had similar survival during the initial stages, but around 24- and 36hpi, the survival of *D. simulans* males is significantly lower (*p*-value < 0.001). Overall, *D. melanogaster* females had markedly higher survival than *D. melanogaster* males and both males and females of *D. simulans* (Figure 2).

### 3.2 Bacterial load and BLUD

Bacterial load determination is a measure of the ability of the immune response to ease bacterial growth rates. The bacterial load data showed no statistical differences among experimental groups when comparing the samples across sexes by each interval for either species (Figure 3, File S14). Previous studies in the females of these two species also detected no significant bacterial load differences upon gram-negative *Serratia marcescens* infection at 12hpi and 25hpi (Hill-Burns and Clark 2010). The data showed that the loads were similar for most individuals in the early stages, at 4- and 12hpi. Overall, the load increased in all the experimental groups for *D. melanogaster* and *D. simulans* at 16- and 20hpi; however, a subset of flies in each group showed a higher load than other individual flies. At 20hpi, both the females and males showed a binary outcome of the infection clustering into two groups. This can be explained if some individuals are able to suppress the bacterial infection progression and keep the bacterial count lower than average, whereas others are less able to inhibit the infection (Duneau *et al*. 2017a).

**Figure 3:**
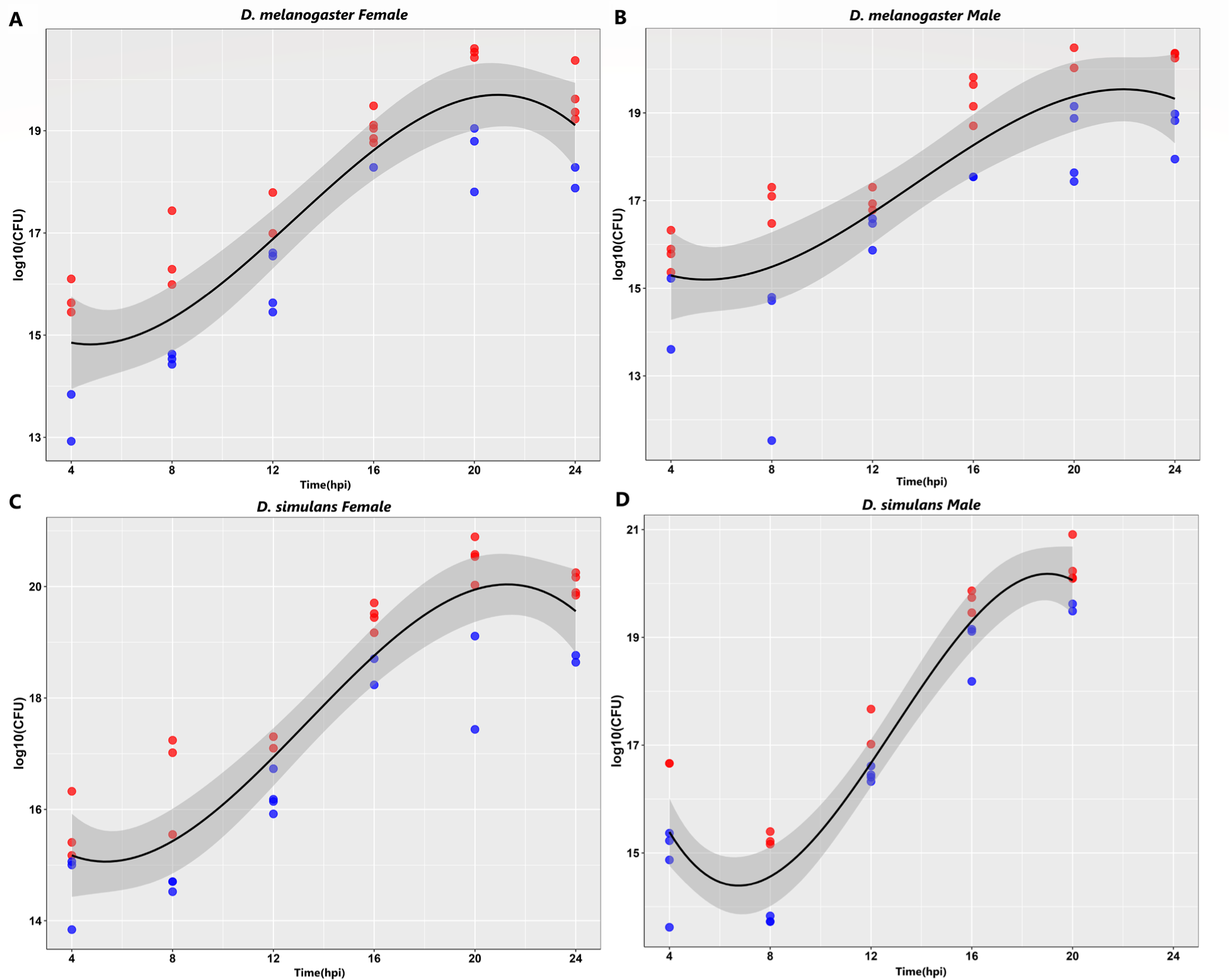
Bacterial load for *D. melanogaster* and *D. simulans* sexes. Determination of bacterial load for females and males at 4-hour intervals upon *P. rettgeri* (PRET) infection in *D. melanogaster* and *D. simulans*. The data were represented for the initial (4, 8, 12hpi) and later (16, 20, 24hpi) stages of the infection in log10(CFU/ML) between the initial and later stages. None of the sample groups showed significant differences at any individual time points (4, 8, 12, 16, 20, and 24hpi) for the bacterial load at *p*-value < 0.05. Polynomial regression was employed to fit a polynomial curve, where bacterial loads above the curve were marked in red and those below in blue. All *D. simulans* males died at 24hpi. There are 6 replicates for each group, except the dmel female at 4 hrs, which has 5 Replicates (one replicate was dropped as an outlier). The wound control data is not shown; it is available in File S14.

The results of the BLUD assay showed no sexually dimorphic pattern in bacterial burden at the time of death upon *P. rettgeri* infection in *D. melanogaster* and *D. simulans*; moreover, there was no significant difference in the species comparison (*p*-value > 0.05) (Figure 4). Overall, the bacterial load was unable to explain the sex differences observed in survival.

**Figure 4:**
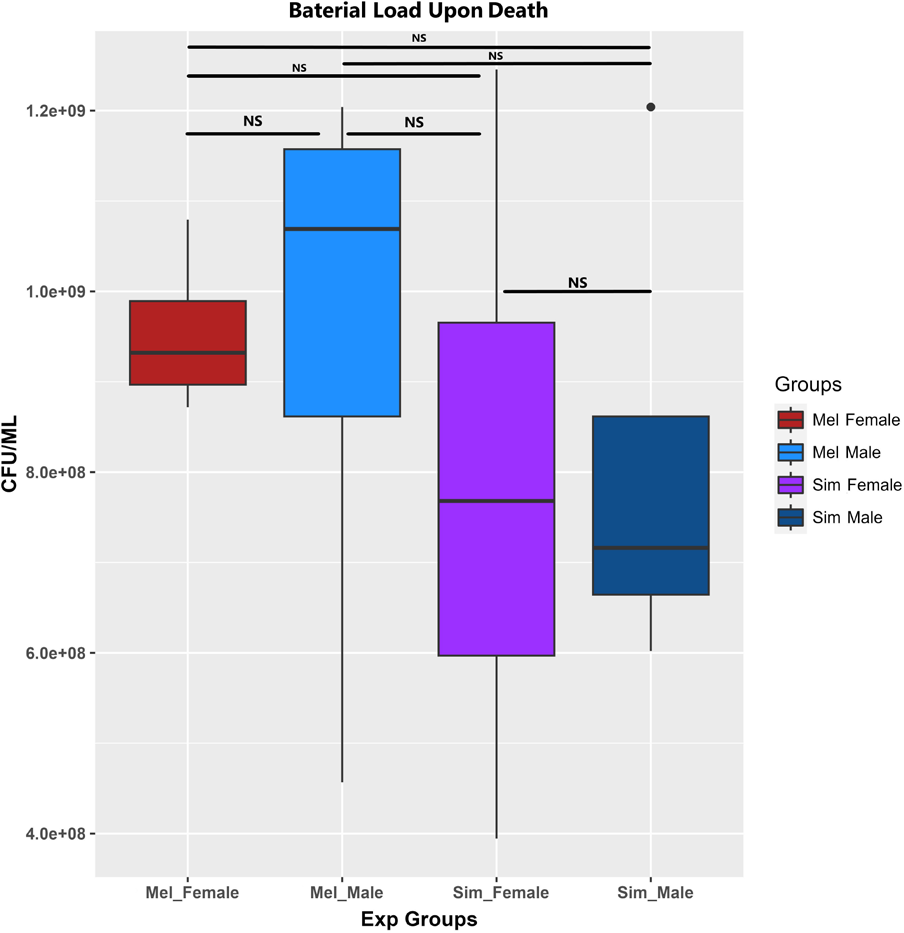
BLUD for *D. melanogaster* and *D. simulans* sexes. Determination of Bacterial Load Upon Death (BLUD) for females and males upon *P. rettgeri* (PRET) infection. The wound control (PBS) showed no death for the experimental groups. The asterisks (*) demonstrated the *p-*values < 0.05, and NS represented no significance.

### 3.3 Identification of DEGs in each sex and species

Within species in *D. melanogaster*, after normalization, 13,588 genes were tested. In response to infection, 2269 genes were downregulated in males compared to 977 in females; with 629 genes downregulated in both sexes (Figure 5A, Table S1). A higher number of genes (1640) were uniquely downregulated in males, compared to females (348), which might indicate higher stress levels in males upon infection. The number of upregulated genes was more comparable between males and females. The number of genes that were upregulated in males was 530, and in females, 599, and 264 upregulated genes were in common (Figure 5A).

**Figure 5:**
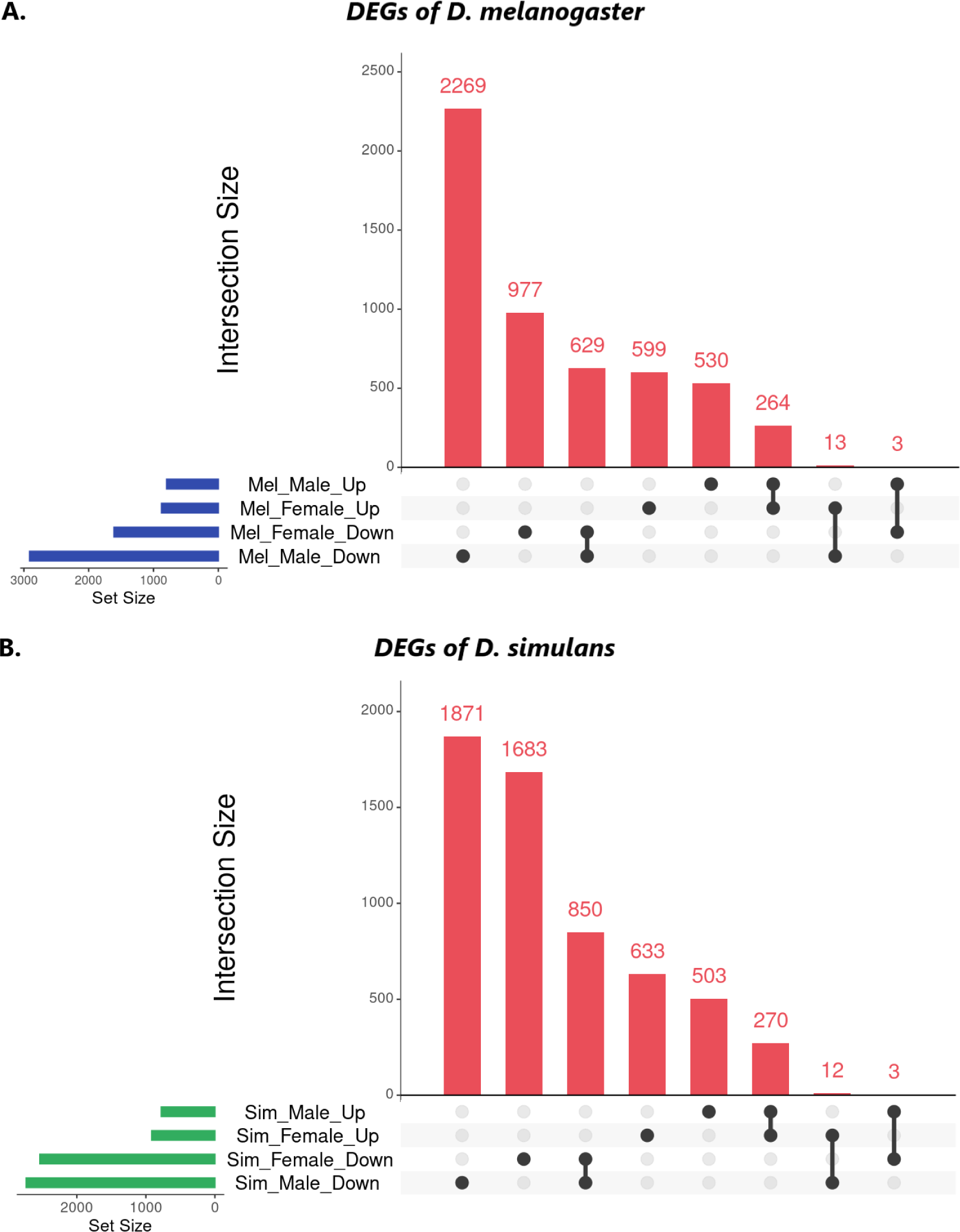
The upset plots represent up- and down-regulated gene counts. **A.** Comparison within species for females and males in *D. melanogaster*, **B.** Comparison within species for females and males in *D. simulans*. The plots also show the synergistically and antagonistically expressed gene numbers for each sex in *D. melanogaster* and *D. simulans*. The figure uses an intersection-based representation (for details, File S4-S5, Table S1).

After normalization, 13,010 genes were tested in *D. simulans,* which also showed a higher number of genes downregulated in males (1871 genes) compared to females (1683 genes), but the difference is less than in *D. melanogaster*, where males had roughly 2-fold higher numbers of downregulated genes relative to females (Figure 5B, Table S1). In females, 633 genes were upregulated, whereas 503 genes increased expression under infection in males. Both sexes downregulated 850 genes and upregulated 270 genes in common in *D. simulans*. The treatment by sex interaction, was tested separately for *D. melanogaster* and *D. simulans* (comparison within species). There were 994 genes (*D. melanogaster*) and 1041 genes (*D. simulans*) that showed sex differences in expression in response to infection with FDR < 0.05 (Figure 5B, Table S1).

To compare across species for each sex, the interaction for the effect of treatment by species was tested separately for females and males (File S6-S7). There were 11,849 genes for which the species could be compared directly. In females, 569 genes were differentially regulated upon infection in the comparison between *D. melanogaster* and *D. simulans* at FDR < 0.05 (File S8). In males, 477 genes have differential regulation in the comparison between these two species (FDR < 0.05) (File S8). Additionally, 159 of the differentially regulated genes are shared between females and males; thus, females showed unique differential regulation for 410 genes and males for 318 genes (File S8).

### 3.4 Comparison of gene expression profiles in female and male adults

The analysis of the DEGs in the experimental groups indicated a complex pattern of sex dimorphism for both species (Figure 6, File S4-S5). However, all experimental groups expressed immune response genes such as AMPs at significantly higher levels than wound control, as expected upon infection. The volcano plots of the four experimental groups showed that *D. melanogaster* females broadly expressed key systemic immune and stress response genes at higher levels than the rest of the three experimental groups. In *D. melanogaster*, both males and females upregulated AMPs, PAMP recognition proteins, regulatory proteins, and stress response genes, but females expressed them at higher levels relative to males. The *D. simulans* females also upregulated immune genes at higher levels than males, but generally, the increase in expression was much lower than that for *D. melanogaster* females (Figure 6).

**Figure 6:**
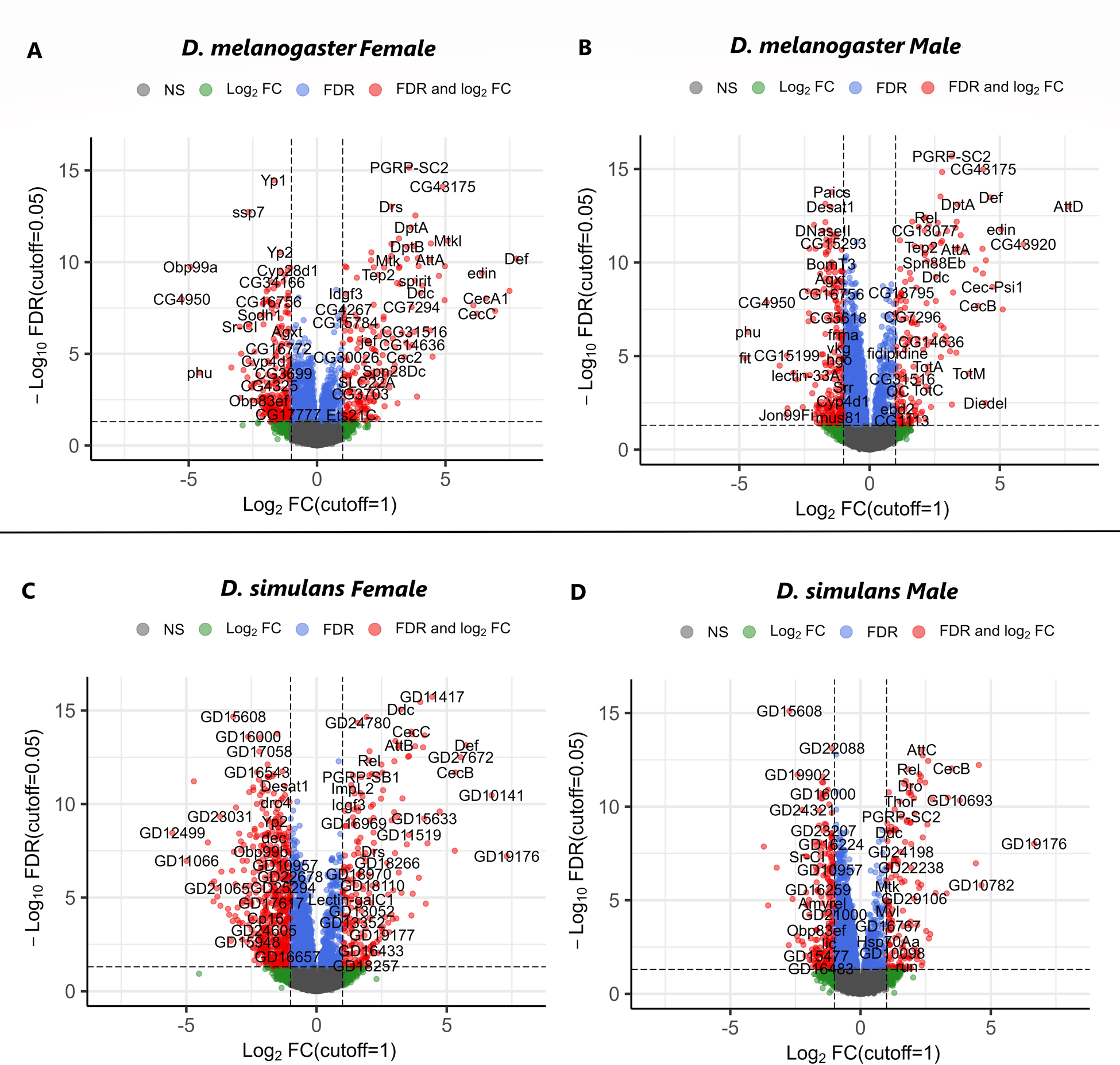
The volcano plots represent DEGs. **A.** *D. melanogaster* females, **B.** *D. melanogaster* males, **C.** *D. simulans* females, and **D.** *D. simulans* males represented upon *P. rettgeri* (PRET) infection compared to wound control (PBS). For *D. melanogaster*, 13588 Normalized Genes, and for *D. simulans*, 13588 Normalized Genes were used to represent DEGs with a cutoff: log2Fold Change ≥ 1; FDR < 0.05. The *D. melanogaster* female also expressed *AttD* (log2FC = 10.52), which is out of the figure range (for details, File S4-S5).

The comparison between species showed that females of the closely related species expressed quite different sets of genes (Figure 6). Comparing across males and females of the species, males suppress different genes than females do in both species. Downregulated genes in females of both species included several female reproductive genes, for example, *Yp1* and *Yp2*. This pattern is explored further in the enrichment analysis. Moreover, as mentioned earlier, the sex differences are visible in the case of the up, and down-regulated gene numbers for females and males in both species; however, the sex differences are more prominent when the data was separated based on the fold changes (FC) (Figure S1, File S4-S5).

### 3.5 Gene set enrichment analysis

Gene set enrichment analysis (GSEA) was conducted using the Gene Ontology Biological Process (GO-BP) annotation. The enriched GO-BP terms showed that males and females expressed defense and stress response gene sets, but they used distinct strategies to different degrees in response to the bacterial infection (File S9).

In *D. melanogaster*, 50 GO-BP terms were significantly enriched in females; among them, 49 were upregulated, and only 1 was downregulated (Figure 7, Table S2, File S9). In males, a total of 101 significantly enriched GO-BP terms were identified, with 84 corresponding to upregulation and 17 to downregulation (Figure 7, Table S2, File S9). Interestingly, 46 out of 49 GO-BP terms for upregulated genes in females are also significant for upregulated genes in males, with 1 out of 1 GO-BP sets of genes commonly suppressed in both sexes.

**Figure 7:**
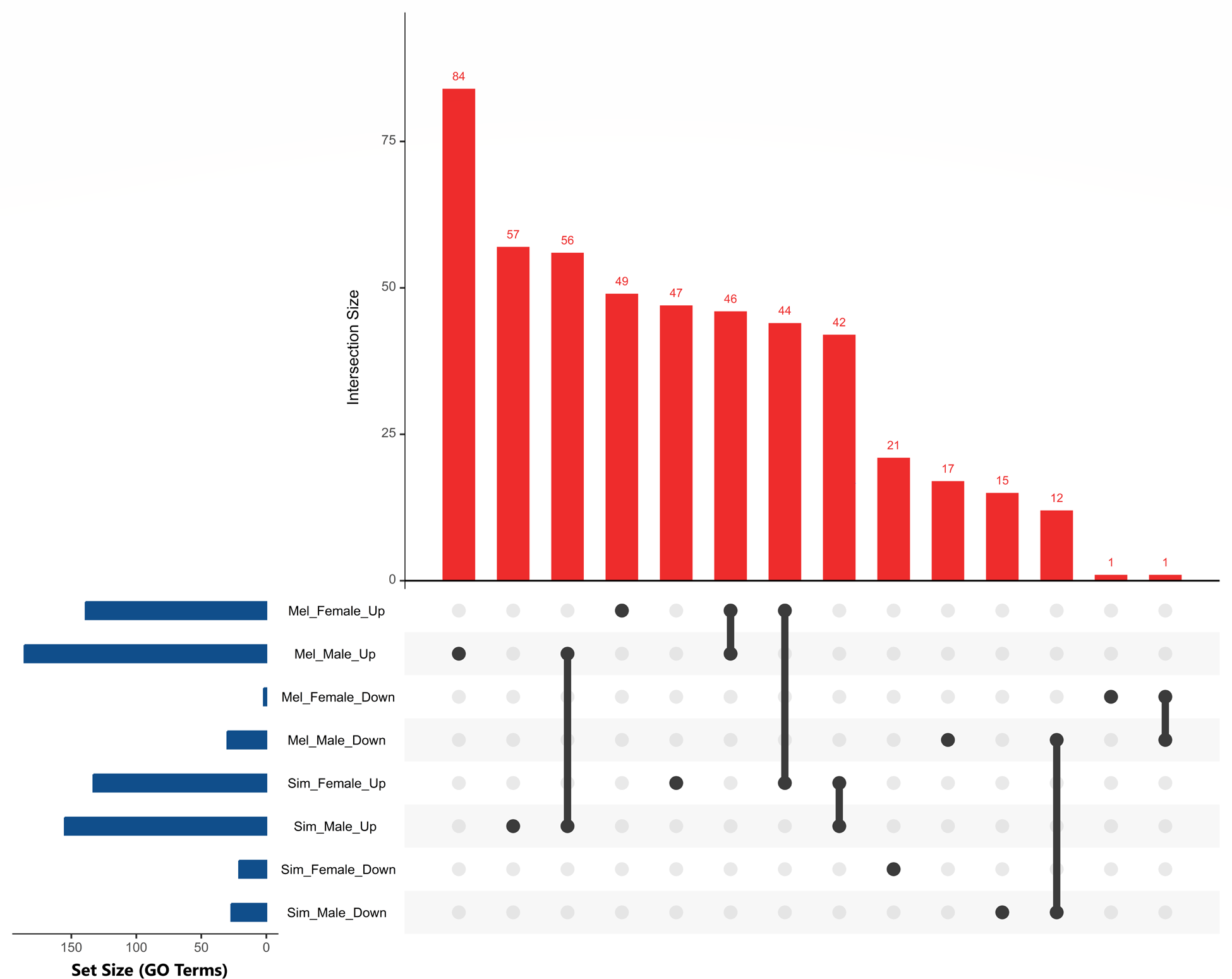
Up- and down-regulated enriched gene ontology terms biological process (GO: BP) in males and females. Mel=*D. melanogaster*, Sim=*D. simulans* (for details, Table S2, File S9). The figure uses an intersection-based representation.

In *D. melanogaster*, the top 20 enriched GO-BP terms for upregulated genes in females showed immune response terms such as defense response and response to gram-negative and -positive bacterium, response to fungus, and antimicrobial humoral response (Figure 8, File S9). *D. melanogaster* males also upregulated gene sets related to similar immune responses, such as response to gram-positive bacterium, antibacterial humoral response, and response to fungus. Females (5 GO-BP terms) and males (7 GO-BP terms) upregulated genes related to negative regulation of the immune and defense responses (Figure 8, File S9). This could correspond to the modulation of immune responses through feedback mechanisms (Tanji and Ip 2005; Vincent and Dionne 2021). The enriched terms corresponding to downregulated genes showed that *D. melanogaster* females downregulate only one GO-BP term, lipid metabolic process; however, in males, 17 GO-BP were identified where almost all of the terms were related to metabolism (14 metabolic processes, 2 catabolic processes, 1 homeostasis) (Figure 8, File S9).

**Figure 8:**
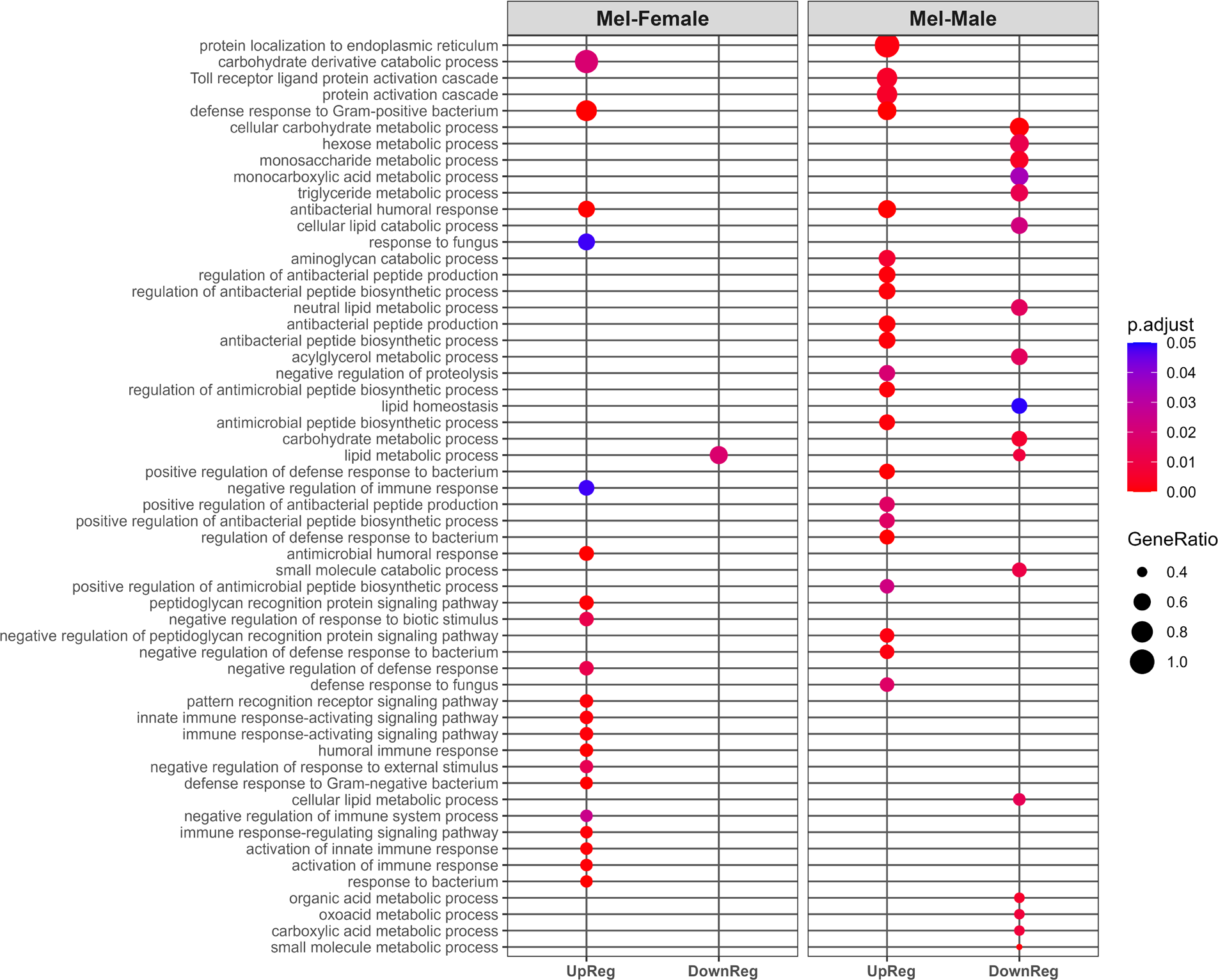
The top 20 up- and down-regulated enriched gene ontology terms (gseGO: BP) in females and males of *D. melanogaster* upon P. rettgeri (PRET) infection (for details, File S9). Mel = *D. melanogaster*, UpReg = Up-Regulation, DownReg = Down-Regulation.

In *D. simulans*, 68 GO-BPs were significantly enriched in females; among them, 47 corresponded to upregulated genes, and 21 downregulated genes (Figure 7, Table S2, File S9). In males, 72 total significantly enriched GO-BP terms were identified, with 57 for upregulated genes and 15 for those that were downregulated (Figure 7, Table S2, File S9). In *D. simulans*, the enriched groups among upregulated genes of the males and females showed 42 commonly activated gene sets, whereas no common GO terms were identified as enriched among downregulated genes (Figure 7, File S9). In *D. simulans*, the top 20 GO-BP terms for the males and females corresponded to activation of the immune response, such as response against gram-negative and positive bacteria, antimicrobial humoral response, innate immune response, etc. (Figure 9, File S9). In addition, both males (6) and females (6) had positively regulated immune-related terms among the top 20 most enriched categories. However, females expressed less terms related to negative regulation of the immune response (3) than males (6) in *D. simulans*. The males of *D. simulans* showed suppression of metabolism-related genes (11 metabolic processes, 4 catabolic processes), which was also observed in *D. melanogaster* males (Figure 8-9). On the other hand, females of *D. simulans* downregulated predominantly reproduction-related gene sets such as eggshell formation, developmental processes involved in reproduction, etc. Thus, although females of both species downregulated the iconic yolk protein genes, widespread downregulation of reproductive genes occurred only in *D. simulans* females (Figure 8-9, File S9).

**Figure 9:**
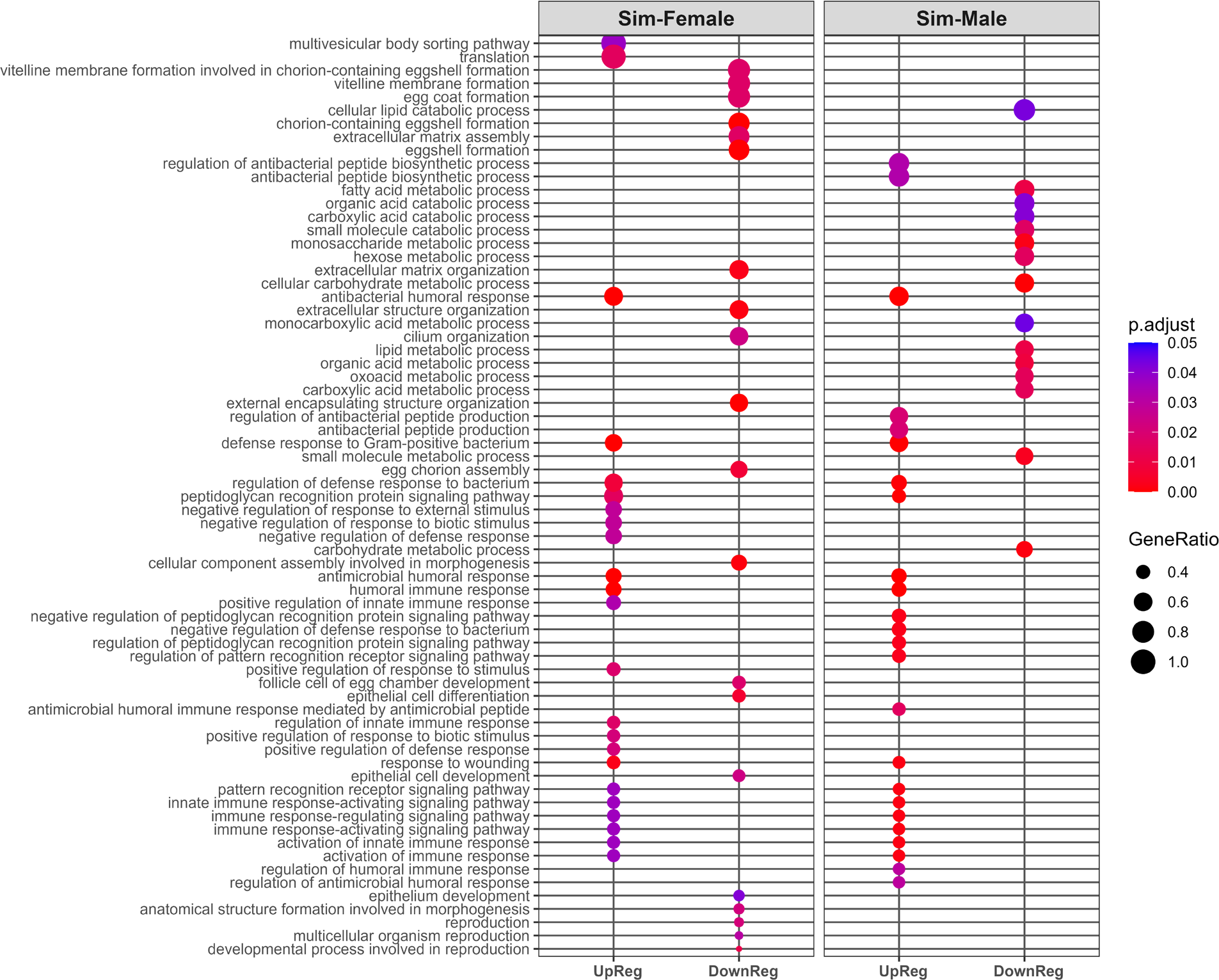
The top 20 up- and down-regulated enriched gene ontology terms (gseGO: BP) in females and males of *D. simulans* upon P. rettgeri (PRET) infection (for details, File S9). Sim = *D. simulans*, UpReg = Up-Regulation, DownReg = Down-Regulation.

In comparisons between species, for females in *D. simulans*, there were 3 upregulated and 21 uniquely downregulated gene sets, whereas for *D. melanogaster* females, there were 5 upregulated terms and 1 downregulated term (File S9). In addition, 44 upregulated terms showed overlap between females of *D. melanogaster* and *D. simulans*, but there is no overlap among the downregulated terms. However, males in both species had 56 upregulated and 12 downregulated GO-BP terms in common. There were 28 uniquely upregulated terms in *D. melanogaster* males and 5 downregulated terms, whereas in *D. simulans* only 1 upregulated term and 3 downregulated terms were unique. Overall, the enrichment analysis showed distinct sex dimorphism with respect to up-or downregulated functional categories of genes in *D. melanogaster* and *D. simulans*. The analysis revealed differences between males and females not only in upregulated biological processes but also in downregulated biological processes, which might reflect different infection levels and control strategies for the sexes (File S9).

### 3.6 Infection-induced differences in the expression of IMD pathway genes

In the IMD pathway, the signal receptors and cellular response showed sex dimorphism in expression in response to infection (Figure 10-11, File S10). This could be caused by differences in the regulation of the IMD pathway between sexes and may be associated with sex differences in the effectiveness of the immune response. In *D. melanogaster*, peptidoglycan recognition proteins (PGRP) of the IMD pathway showed significantly higher expression in females for PGRP-LB and PGRP-LC than in males upon infection (*p*-adj < 0.05) (Figure 10, File S10). The signal transducers for the IMD pathways were not significantly different between sexes; however, the key IMD transcription factor Relish was significantly upregulated more in males (log2FC = 2.16) than in females (log2FC = 1.54) in *D. melanogaster* (*p*-adj < 0.05). The effector genes of IMD showed broad upregulation in both sexes, which includes AMPs such as *Attacins* (*Att*), *Cecropins* (*Cec*), and *Diptericin-B* (*DptB*). In both sexes, *AttD* was the most highly expressed AMP; it was upregulated around 20-fold in females, as compared to 15-fold in males (*p*-adj < 0.05) (File S10). In addition, *CecB*, *CecA1*, *AttB*, *AttA*, and *DptB* were upregulated at significantly higher levels in females relative to males (*p*-adj < 0.05) (File S10). Moreover, females exhibited upregulation in *Dro* (log2FC = 2.89), *DptA* (log2FC = 3.66), *CecC* (log2FC = 6.22), *CecA2* (log2FC = 6.08), and *AttC* (log2FC = 3.78), and these genes are also strongly upregulated in males, *Dro* (log2FC = 2.63), *DptA* (log2FC = 3.38), *CecC* (log2FC = 4.34), *CecA2* (log2FC = 4.46), and *AttC* (log2FC = 3.29) (File S10).

**Figure 10:**
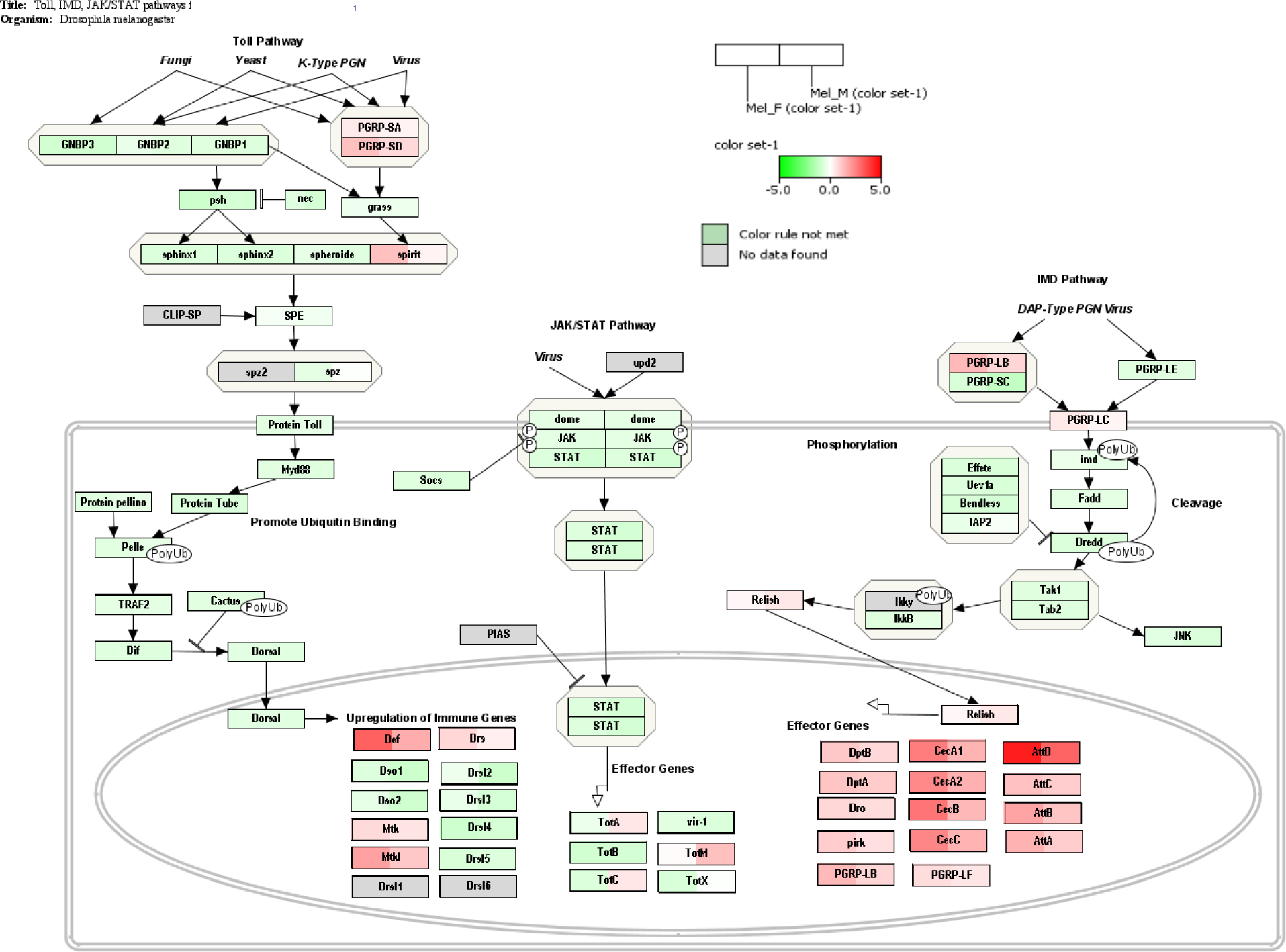
DEGs in females and males of *D. melanogaster* in response to *P. rettgeri* infection for genes in the Toll-IMD-Jak/Stat signaling pathway (wiki pathway ID: WP3830). The expression (z-score) of the pathway is represented without statistical cut-off. Red represents upregulation, green represents downregulation, and white represents no change compared to wound control (for details, File S10, Figure S2).

**Figure 11:**
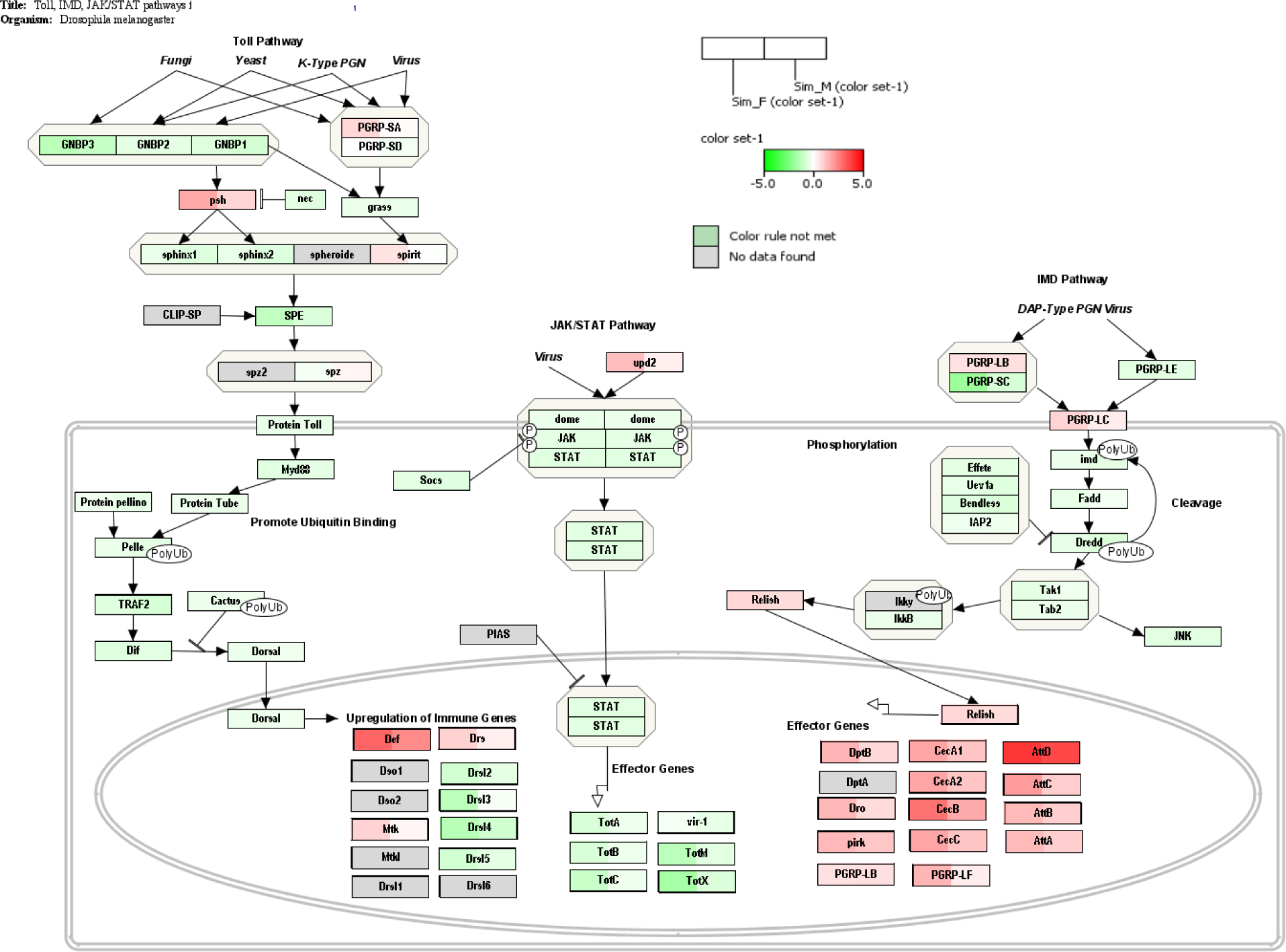
DEGs in females and males of *D. simulans* in response to *P. rettgeri* infection for genes in the Toll-IMD-Jak/Stat signaling pathway (wiki pathway ID: WP3830). The expression (z-score) of the pathway was represented without statistical cut-off. Red represents upregulation, green represents downregulation, and white represents no change compared to wound control (for details, File S10, Figure S2).

In *D. simulans*, the IMD pathway also showed female-biased upregulation in response to bacterial infection, but not all sex differences in specific genes are conserved between species (Figure 11, Figure S2, File S10). The focal signal-transducing innate immune signal receptor PGRP-LC is expressed significantly higher in females relative to males in *D. simulans* when each group is compared with its wound control (*p*-adj < 0.05) (Choe *et al*. 2005). Interestingly, PGRP-LF, which inhibits the activation of IMD pathways by capturing circulating peptidoglycan, is highly upregulated in *D. simulans* females (log2FC = 3.08) compared to the other three experimental groups, and also showed significant sex differences with *D. simulans* males (log2FC = 1.34) (*p*-adj < 0.05) (Figure 11, File S10) (Maillet *et al*. 2008). The key IMD transcription factor Relish was upregulated in females (log2FC = 2.02) and males (log2FC = 1.84) in *D. simulans*. The effector genes of IMD showed broad upregulation in both sexes, which includes AMPs such as *Attacins* (*Att*), *Cecropins* (*Cec*), and *Diptericin-B* (*DptB*). In both sexes, *AttD* was the most highly expressed AMP; it was upregulated around 14.6-fold in females, as compared to 13.3-fold in males (*p*-adj < 0.05) (File S10). In addition, *AttA*, *AttB*, *AttC*, *CecA1*, *CecA2*, *CecB*, *CecC*, *Dro*, *DptA*, and *DptB* were upregulated at significantly higher levels in females relative to males (*p*-adj < 0.05) (File S10). Overall, the IMD cellular effector molecules of *D. simulans* showed higher expression in females than males. However, the degree of upregulation of these AMPs in *D. simulans* females is lower on average than in *D. melanogaster* females (File S10).

### 3.7 Regulators of IMD pathway genes

The positive and negative regulators of the IMD pathway were analyzed for each sex in both species. In the case of the IMD positive regulators (FBgg0001197), PGRP-SD, which can significantly contribute as a ‘positive regulator’ of the IMD pathway, showed significant differential expression between females (7.64-Fold) and males (5.56-Fold) of *D. melanogaster* (*p*-adj < 0.05) (Figure S3, File S11) (Gelbart *et al*. 1996). Expression patterns of this gene also differed between females and males in *D. simulans* (*p*-adj < 0.05); however, both sexes of *D. simulans* showed lower levels of upregulation than their *D. melanogaster* counterparts. In addition, expression of the gene encoding Charon, which can directly activate Rel/NF-kB, was upregulated in males (log2FC = 1.19) to a higher degree than in females (log2FC = 0.36) in only *D. melanogaster* (*p*-adj < 0.05). Differences in the ‘negative regulators’ of IMD (FBgg0001196) also included significantly higher expression of PGRP-SC2, a down-regulator of the IMD pathway, and PGRP-LB in *D. melanogaster* females relative to males (*p*-adj < 0.05; Figure S3, File S11) (Bischoff et al. 2006; Guo et al. 2014). While PGRP-LB is one of the prominent receptors involved in the recognition and activation of the IMD pathway, researchers have found that it enables females to reduce IMD activity upon a decrease in bacterial numbers (Zaidman-Rémy et al. 2006). Thus, the significant upregulation of two negative regulators of IMD, PGRP-LB and PGRP-SC2, in *D. melanogaster* females tempts us to speculate that in females, there is more controlled IMD expression relative to males (Bischoff et al. 2006; Zaidman-Rémy et al. 2006; Guo et al. 2014). Additionally, the negative regulator of IMD, PGRP-LB and PGRP-SC2, were not highly upregulated in *D. simulans* compared to *D. melanogaster* sexes (*p*-adj < 0.05). However, another negative regulator, PGRP-LF, is highly upregulated in *D. simulans* females (log2FC = 3.08) compared to males (log2FC = 1.34) (*p*-adj < 0.05) (Figure S3) (Maillet et al. 2008). Overall, the IMD pathway is involved in the response to *P. rettgeri* via effector molecules for females and males in a sex differential manner in *D. melanogaster* and *D. simulans*; however, higher upregulation of the negative regulators such as PGRP-LB, PGRP-SC2, and PGRP-LF in females might indicate that these flies activated IMD pathways at a level where suppression was initiated, whereas other experimental groups did not reach that elevated level.

### 3.8 Infection-induced differences in the expression of Toll pathway genes

The Toll pathway was originally identified as an effector pathway activated upon gram-positive bacterial and fungal infection, but recent studies show it also plays a crucial role in the responses to gram-negative bacterial infections and viral infections (Valanne et al. 2022). In *D. melanogaster*, signal receptor PGRP-SD was more elevated in females than males (*p*-adj < 0.05) (Figure 10, Figure S2, File S10). In addition, the extracellular molecule *Serine Protease Immune Response Integrator* (*spirit*) showed significant differences between females (log2FC = 3.81) and males (log2FC = 1.73) (*p*-adj < 0.05). The males significantly upregulated *spz;* Spz is a core ligand of Toll receptor (Tl), relative to females in *D. melanogaster* (*p*-adj <0.05) (Valanne et al. 2011). The ‘intracellular signal transducers’ of Toll had no significant changes in regulation between sexes, including the key NF-κB-related transcription factors Dorsal-related immunity factor (Dif) and Dorsal (dl) (Valanne et al. 2011). In contrast, the effector molecules showed higher upregulation of *Drosomycin* (*Drs*), *Defensin* (*Def*), and *Metchnikowin*-*like* (*Mtkl*) in females relative to males in *D. melanogaster* (*p*-adj < 0.05). *Mtkl* was the most highly expressed Toll-AMP for both sexes in *D. melanogaster*, which was upregulated by approximately 10-fold in females and 7-fold in males (*p*-adj < 0.05). Interestingly, the gene encoding focal antifungal peptide-producing AMP, *Drosomycin* (*Drs*), was also upregulated in both sexes, but other antifungal AMP genes such as *Daishos* (*Dso1* and *Dso2*) and *Drosomycin*-*like* (*Drs1*-*6*) showed no significant increase in expression upon bacterial infection, instead some of them including *Dso1* and *Dso2* showed significant downregulation, where these genes were downregulated in males to a higher degree than in females (*p*-adj < 0.05) (Figure 10, File S10).

*D. simulans* showed analogous expression patterns to *D. melanogaster* in the extracellular portion of the Toll Pathway. PGRP-SA and PGRP-SD were significantly upregulated at higher levels in females relative to males (*p*-adj < 0.05), but the degree of upregulation is lower than in *D. melanogaster* (Figure 10-11, File S10). This also holds true for the expression of *spirit*, which exhibits significant sex differences (*p*-adj < 0.05), although the expression level is lower compared to *D. melanogaster*. The gene *spz* was significantly overexpressed in males relative to females (*p*-adj < 0.05), as also identified in *D. melanogaster* (Figure 10-11). In *D. simulans*, the key NF-κB-related transcription factors *Dorsal-related immunity factor* (*Dif*) and *Dorsal* (*dl*) showed no significant differences between sexes (Valanne et al. 2011). The effector molecules showed higher upregulation of *Drosomycin* (*Drs*), *Defensin* (*Def*), and *Metchnikowin* (*Mtk*) in females relative to males In *D. simulans* (*p*-adj < 0.05). Interestingly, *Drs* was also upregulated in both sexes, but other antifungal AMPs such as *Drosomycin*-*like* (*Drsl*) *2*, *Drsl 4*, and *Drsl 5* showed significant downregulation upon bacterial infection, where females showed higher levels of downregulation than males did (*p*-adj < 0.05) (Figure 10, File S10).

In between species comparisons, *D. simulans* showed significant downregulation of *SPE*, unlike *D. melanogaster*, with females (log2FC = −1.3) exhibiting more downregulation than males (log2FC = −0.44) (*p*-adj < 0.05). Another key difference was observed in the expression of Clip-SP *Persephone (psh)*, *D. simulans* females (7-Fold) and males (4-Fold) upregulated *psh*, where the reverse direction is observed in females (log2FC = −0.81) and males (log2FC = −0.45) of *D. melanogaster*. Overall, these findings suggest that *D. simulans* might be using a different route to stimulate Toll pathways other than PRR base activation.

### 3.9 Regulators of Toll pathway genes

The regulation of the Toll pathway is much more complex than the regulation of IMD, as there are diverse functions performed by the Toll pathway in addition to immunity. Drosophila possesses six distinct Toll receptors, each contributing individually or in concert to accomplish complex multidimensional functions; the data showed none of the Toll receptors were sex differentially expressed in either species (Valanne et al. 2011) (File S11). However, the *spz* activation pathway, which is critical to the regulation of the Toll pathway as Spz can be activated in multiple ways, such as PRR-based, Psh-SPE-Spz, and Psh-Protease-Spz, shows apparent sex differences. Multiple genes in this pathway are expressed in a sex differential manner, such as *spirit, psh*, *Hayan*, and *PGRP-SC2* (Figure S4, File S11). The sex differential regulation of *spirit* and *psh* has previously been discussed with respect to the overall Toll pathways (Duneau et al. 2017b). There were also female-biased sex differences for wound-related serine protease Hayan in *D. simulans* (*p*-adj < 0.05), which is involved in the activation of the melanization cascade as well as Spz based Toll pathway activation (Yamamoto-Hino and Goto 2016; Dudzic et al. 2019). *Hayan* was also upregulated in females (log2FC = 3.22) and in males (log2FC = 2.45) in *D. melanogaster*. The gene encoding PGRP-SC2, which acts as a negative regulator of the IMD pathway, is expressed at higher levels upon infection in females (log2FC = 3.59) than males (log2FC = 3.15) in *D. melanogaster* (*p*-adj < 0.05) (File S12) (Bischoff et al. 2006). *PGRP-SC2* was also upregulated in both females (log2FC = 1.60) and males (log2FC = 1.56) in *D. simulans*. (File S12). Aside from the *spz* activating pathway, positive (FBgg0001057) and negative (FBgg0001053) regulators of the Toll pathway showed no significant sex differences in either species (Figure S4, File S11).

Overall, considering the Toll pathway and its regulators, suggests there is female-biased sex differential upregulation of Toll-AMPs for both species; however, the degree of upregulation is higher in *D. melanogaster* than in *D. simulans* like IMD-AMPs. A comparison between species detected a remarkable divergence in the Toll activation process, where *D. simulans* employed both PRR-dependent and independent mechanisms, while the closely related *D. melanogaster* solely upregulated genes related to PRR-dependent activation.

### 3.10 Other immune response molecules and pathways

The innate immune response of Drosophila is a complex interplay of conserved innate immune signaling pathways and other immune molecules, along with the involvement of immune cells. Several Drosophila immune-induced molecules (DIMs) have been identified in the fly hemolymph (Uttenweiler-Joseph et al. 1998). Recent research revealed that twelve related immune-induced molecules, known as the Bomanins (Boms), constitute a family that plays a role in defending against certain fungi and bacteria (Clemmons et al. 2015). Three of the twelve Bomanins (Boms) in *D. melanogaster*—*BomT1*, *BomS5*, and *BomBc3*—are expressed in a sex-differential manner (*p*-adj < 0.05). (Figure S5, File S11). *BomT1* is upregulated to a greater degree in females (log2FC = 2.59) relative to males (log2FC = 1.53) (*p*-adj < 0.05). Opposing sex differences were observed for *BomS5*, where females upregulated, and males downregulated expression (*p*-adj < 0.05). *BomBc3* was downregulated in both sexes in *D. melanogaster*, but in males, it was downregulated to a greater extent than in females (*p*-adj < 0.05). However, only *BomS2* and *BomS4* showed significant sex differences in *D. simulans* (*p*-adj < 0.05). In the case of *BomT1*, upregulation was observed in both females (log2FC = 1.18) and males (log2FC = 1.03) in *D. simulans*. *BomBc3* expression is different from that of *D. melanogaster,* where *BomBc3* is downregulated in females (log2FC = −1.36) and males (log2FC = −1.10) (Figure S5, File S11). The expression of the Boms and their regulators showed marked species differences among the sexes.

Melanization is one of the extraordinary features of insect innate immunity and is regulated by Toll pathways in terms of the gene expression of serine proteases (SPs) and serpins (Spn) (Dudzic et al. 2019). PGRP-SA and GNBP1/ β-Glucan Recognition Proteins (βGRP) can directly modulate the expression of the melanization cascade genes (Matskevich et al. 2010). In the case of core serine proteases, *Hayan* and *SP7*/*MP3* both show significant sex differential expression in *D. simulans* (*p*-adj < 0.05) (Figure S5, File S11). Although *D. melanogaster* females and males upregulated Hayan and SP7 more than wound control, no significant sex differences were observed. In both species, the level of *SP7* upregulation is lower than that of *Hayan*. Although key SPs are upregulated, there is downregulation of the key *prophenoloxidases* (*PPO*), *PPO1*, and *PPO2* (Figure S5, File S11). In addition, there is significant upregulation of the negative regulator of melanization, *Spn28Dc*, in both species. With higher upregulation in females (log2FC = 2.99) relative to males (log2FC = 1.15) in *D. melanogaster* (*p*-adj < 0.05) and a similar pattern found in the females (log2FC = 3.78) and males (log2FC = 2.07) of *D. simulans* (*p*-adj < 0.05) (Figure S5, File S11). These data strongly suggest that the melanization cascade was not activated; thus, Hayan and SP7 might be utilized by other immunological processes, such as Toll activation, as has been observed in other studies (Yamamoto-Hino and Goto 2016; Dudzic et al. 2019).

In *Drosophila*, the Jak-STAT pathway is crucial for regulating antimicrobial peptide (AMP) production in response to bacterial infection and antiviral response (Myllymäki & Rämet, 2014). The cytokine-like ligands *upds*, signal receptor *domeless* (*Dome*), *Janus kinase* (*JAK*), *hopscotch (hop)*, and the Drosophila STAT transcription factor, *stat92E*, were not sex-differentially expressed in either species (Figure S7, File S10-11). Despite the lack of significant sex differences in transcript level in upstream parts of this pathway, some of the effector molecules, such as *Turandot C* (*TotC*), *TotA*, *TotX*, and *TotM*, were significantly upregulated compared to the wound control in males of *D. melanogaster (p-*adj < 0.001); this pattern is absent in females of *D. melanogaster* and both sexes of *D. simulans* (Figure 10-11). The study found no sex differences in components of negative regulators in either species (Figure S7, File S11). JNK is also known as the TNFalpha-Eiger pathway in Drosophila, which, like Jak-STAT, has a diverse set of functions with a key role in immune responses, development, and morphogenesis (Gan et al. 2021). The gene expression analysis found that JNK receptors, protein kinase, and regulators showed no significant sex differences (Tafesh-Edwards and Eleftherianos 2020)(Figure S7, File S11).

Finally, apart from PGRPs of the IMD and Toll pathways, our data found *PGRP-SB1*, an antibacterial PGRP, was more upregulated in females (log2FC = 2.90) relative to males (log2FC = 2.11) in *D. melanogaster* (*p*-adj < 0.05) (File S11). The upregulation is also conserved in *D. simulans*, where females (log2FC = 1.71) upregulate *PGRP-SB1* more than males (log2FC = 0.84) (*p*-adj < 0.05) (File S11). Prior works on *PGRP-SB1* report that it can be significantly upregulated upon IMD pathway activation by the gram-negative bacterial infection, which is consistent with our findings (Zaidman-Rémy et al. 2011). The analysis of *thioester-containing proteins* (*Teps*) identified that only *Tep2* showed differential expression compared to wound control in females (log2FC = 2.38) and males (log2FC = 1.98) in *D. melanogaster* (Figure S6, File S11). The females (log2FC = 0.44) and males (log2FC = 0.32) also upregulated *Tep2* in *D. simulans*, but the degree of upregulation is significantly lower than in *D. melanogaster* counterparts (Figure S6, File S11).

## 4. Discussion

The emergence of sex differences in immune responses is thought to originate from complex evolutionary dynamics including trade-offs and sexual conflict (Zuk and Stoehr 2002; Scharf et al. 2013; Schwenke et al. 2016). The interplay between the immune system and various physiological processes can result in trade-offs between immunity and other energetically costly life-history traits, such as reproduction (Zuk and Stoehr 2002; Schwenke et al. 2016). Immune responses can also be sexually dimorphic, with males and females showing different responses to the same pathogens in identical environments (Lazzaro and Little 2009; Morrow 2015). Recently, there has been a growing interest in adult immunity and the role of sex-biased gene expression in immune response pathways in generating sex dimorphism in the overall immune response due to its impact on the medical sector (Fischer et al. 2015; Wilkinson et al. 2022). However, many studies in immunology have focused predominantly on one sex, often males (Belmonte et al. 2020). The importance of including both sexes in immunological studies is increasingly acknowledged and enables researchers to achieve a comprehensive understanding of sex dimorphism, evolution, and molecular mechanisms of the immune response.

Our investigation of sex differences in immune response against *P. rettgeri* in *D. melanogaster* and *D. simulans* found that survival differences may not be conserved, and sexes differ in survival in *D. melanogaster*. Interestingly, we also find that although both sexes express similar immune response genes for conserved immune pathways and some other immune response components, there is a strong pattern of a robust response in *D. melanogaster* females relative to males. In *D. simulans*, females also mounted higher immune gene expression, but the expression difference between the two sexes is not as high as in *D. melanogaster* and did not confer a significant survival advantag*e*. Moreover, both species upregulate similar genes to inhibit infection, but broadly, the degree of up or downregulation is lower in *D. simulans* than in *D. melanogaster*. Overall, these findings highlight species-specific variations in the effectiveness of the immune response and the physiological impact of the infection. In addition, the downregulation of metabolism-related gene sets in males of both species, and the downregulation of reproductive genes in *D. simulans* suggests that each sex uses different strategies to modulate other processes in response to the infection. Lastly, the evolutionarily conserved bacterial pathways IMD and Toll exhibited sex-specific variation, encompassing not only effector molecules but also regulators, such as those involved in negative feedback, in the context of infection progression. This suggests that the IMD pathway responds more in a controlled manner in *D. melanogaster* females than in the rest of the three groups. Moreover, in *D. simulans,* infected animals used two distinct strategies at sixteen hours post-infection, canonical PRR-SPE-Spz, and microbial proteases-based Persephone cascade, to activate the Toll pathways.

### 4.1 Survival of unmated males and females differs between the sexes and species

The survival rates of unmated male and female flies were monitored over a 36-hour period following infection. In *D. melanogaster*, females showed significantly higher survival rates than males from 18hpi onward. In *D. simulans*, while females had slightly higher mean survival rates than males, the difference was not statistically significant. In previous studies, females have been observed to represent higher survival rates than males, for example, in gram-positive *S. aureus* infection in *D. melanogaster* (Duneau et al. 2017b). However, this same study also found that *D. melanogaster* females were more susceptible to *P. rettgeri* (gram-negative), *P. alcalifaciens* (gram-negative), and *E. faecalis* (gram-positive) infections (Duneau et al. 2017b). In addition, two individual studies of gram-negative *Pseudomonas aeruginosa* infections showed opposite-sex survival bias direction (Chowdhury et al., 2019; Vincent & Sharp, 2014).

Sex dimorphism in survival rates appears to depend on sex, host strains, type of infection, and external conditions in *Drosophila* (Howick and Lazzaro 2014; Rai et al. 2023). It has also been shown that even when the pathogen and host are identical, mating status, diet, age, and injection site, among other differences, can alter the outcome of the infection (Lazzaro and Little 2009; Short and Lazzaro 2010; Howick and Lazzaro 2014; Rai et al. 2023). Therefore, results from different studies may vary due to differences between laboratories or due to specific differences in the study design.

In our study, both males and females were unmated, whereas in the Duneau et al. 2017 study, males were mated, and then compared to mated or unmated females (Duneau et al. 2017b). Studies have indicated that mated females exhibit increased susceptibility to infection, elevated pathogen loads, and reduced capacity to initiate immune responses, which could explain why unmated females in our study had better outcomes than in studies that focused on mated animals (Short and Lazzaro 2010; Schwenke and Lazzaro 2017; Rai et al. 2023).

To the best of our knowledge, the post-infection survival of *D. simulans* has not been previously reported; thus, a direct comparison to existing studies is not possible. However, when comparing between species, *D. melanogaster* females had significantly higher survival rates relative to *D. simulans* females after the infection was established. Similarly, *D. melanogaster* males showed significant differences in survival rates compared to *D. simulans* males from 18hpi. Overall, there are both sex- and species-based differences in survival rates, with *D. melanogaster* females showing the highest survival rates. Previous studies have shown variation within the same species based on different conditions, as our results also show compared to prior studies. Here, we have different species in the same environment and with concurrent experiments where both sexes and species can be directly compared. This means that the difference we observe between *D. melanogaster* and *D. simulans* is likely to be genetic, although we note that evaluation of additional strains is needed to establish fixed species-level differences. This study showed that the impact of sex on survival may have diverged between these species and both sex and species effects may be contributing to differences observed.

### 4.2 Distinct patterns of up and down gene regulation are concordant with differences in survival

The up and downregulation of gene expression in the context of stress is a common phenomenon (Lecheta et al. 2020; Girgenti et al. 2021). There are fewer upregulated genes than downregulated for both females and males in both species. *D. melanogaster* females downregulated 977 genes compared to 2269 in males, which is consistent with lower survival rates of males and may indicate higher stress conditions in males relative to females. Compared to the 1871 genes downregulated in *D. simulans* males and 1683 genes in females, *D. melanogaster* females downregulated fewer genes. A similar pattern of up and down-regulation was observed upon Sigma Virus infection in *D. melanogaster*, where males downregulated 336 genes compared to 88 genes in females (Carpenter et al. 2009).

The enrichment analysis indicated differences between sexes in which types of genes were downregulated in *D. melanogaster*. Only one GO-BP category was significantly enriched among downregulated genes in females, whereas 17 metabolism-related downregulated categories were identified in males. This is consistent with the idea of higher stress levels in infected males relative to females in *D. melanogaster*. Likewise, *D. simulans* males downregulated 15 GO-BP categories, among which metabolism-related genes were predominant.

Previous studies also found a trade-off between immune and metabolic processes in males upon bacterial infection in *D. melanogaster* (Schlamp et al. 2021). Schlamp et al. showed that the activation of the IMD pathways affects the metabolic homeostasis of *Drosophila* males (Schlamp et al. 2021). *D. simulans* females may also have been under increased levels of stress because of infection, given that there was extensive downregulation of reproductive GO: BP terms, such as eggshell formation and reproductive developmental processes. Multiple studies have reported trade-offs between immune response and reproduction in both males and females in *Drosophila* as well as other insects (McKean and Nunney 2001; Short et al. 2012; Schwenke et al. 2016). The previous research on *Drosophila* infected with *P. rettgeri* revealed that females halt egg-laying during the acute phase of infection, with a return to normalcy observed during the chronic phase (Howick and Lazzaro 2014) and upon DNA virus Kalithea (KV) infection, females experience degeneration of ovarian tissue and reduced fecundity (Palmer et al. 2018). Additionally, viral-induced female infertility was found upon flock house virus (FHV) infection due to the destruction of oocytes (Thomson et al. 2012). Overall, *D. melanogaster* females downregulated fewer genes, and had fewer functional categories that were consistently downregulated, while at the same time upregulating immune-related genes to a greater degree, which may be related to their higher survival rates.

### 4.3 Sex and species differences in the IMD pathway

IMD receptors and effector molecules were upregulated in females and males in both species. In general, females showed higher levels of expression than males, especially the effector molecules such as AMPs. There is a noticeable degree of higher upregulation in *D. melanogaster* flies compared to *D. simulans*. As the IMD pathway is specialized in response to gram-negative bacterial infection, *P. rettgeri* infection is expected to elicit a higher level of IMD-regulated AMP expression than other core immune response pathways. There was a consistent pattern of female-biased sex differential expression in both species for IMD-AMPs; however, it was also evident that the sexes of *D. simulans* consistently showed lower levels of upregulation than *D. melanogaster* sexes.

The species-level expression difference is also visible in the positive and negative regulators of the IMD pathway; for instance, PGRP-SD, primarily associated with the Toll pathway, significantly contributes to the positive regulation of the IMD pathway by amplifying the recognition of peptidoglycans (Tanji and Ip 2005; Myllymäki et al. 2014). Our results showed that *D. melanogaster* males significantly upregulate the gene *Charon,* which can directly activate the NF-kB/Relish at AMP promoters in males via a PARP-1/Relish/Charon-dependent innate immune response (Ji et al. 2016). Alternatively, the knockdown of *Charon* expression plays a vital role in the translocation of Relish to the nucleoplasm, and blocks that function making flies vulnerable to bacterial attack (Ji et al. 2016). The influence of the PARP-1/Relish/Charon-dependent response might act in conjunction with greater upregulation of NF-kB/Relish in *D. melanogaster* males, relative to females; males may amplify the response to compensate for lower upregulation of signal receptors (PRRs) (File S11). This indicates that sex differences are likely to be complex, and interactions between different genes and pathways will need to be untangled to delineate key mechanisms (Ji et al., 2016).

The precise modulation of energetically demanding immune pathways is crucial to halt the progression of infection. Over-activation of the IMD pathways can severely affect survival (Bischoff et al. 2006). A comprehensive analysis of the negative regulators of IMD pathways was conducted, revealing that *D. melanogaster* and *D. simulans* exhibited sex differences in expression of the genes encoding certain proteins, namely PGRP-LB, PGRP-SC2, PGRP-LF, and Pirk, which play roles in the negative regulation of IMD pathways. *D. melanogaster* females and males both upregulated PGRP-LB, PGRP-SC2, PGRP-LF, and Pirk, but differed in the degree of upregulation. PGRP-LB works as an activator of the IMD pathway, but excessive upregulation of PGRP-LB can trigger a negative feedback loop for the IMD pathway (Bischoff et al. 2006; Vincent and Dionne 2021). *D. melanogaster* females significantly upregulated PGRP-LB more than males, and both sexes of *D. simulans. D. melanogaster* females also upregulated IMD-regulated AMPs at a significantly higher level than the rest of the groups. The significant overexpression of PGRP-LB could either activate the IMD pathway to more efficiently eliminate pathogens, or lead to the pathway’s downregulation via negative feedback in *D. melanogaster* females (Figure S2, File S11).

In addition to PGRP-LB, PGRP-SC2, another negative regulator of IMD pathways, also showed significantly higher upregulation in *D. melanogaster* females relative to others. Another member of the PGRP, PGRP-LF, is a membrane-associated protein, but it cannot bind to peptidoglycans directly (Persson et al. 2007). PGRP-LF interacts with PGRP-LC and functions as a negative regulator of IMD signaling (Maillet et al. 2008; Basbous et al. 2011). Both females (log2FC = 2.61) and males (log2FC =2.27) upregulated PGRP-LF higher than control in *D. melanogaster*, with no significant difference between the sexes. Interestingly, PGRP-LF is highly upregulated in *D. simulans* females compared to the other three groups (Maillet et al. 2008). Another PGRP-LC interacting protein, Pirk, which is involved in the negative regulation of IMD, was upregulated at significantly higher levels in females and males of both species (Vincent and Dionne 2021).

Overall, the IMD pathway was employed in a sex-differential manner in *D. melanogaster* and *D. simulans*. However, upon comparing these four “brakes” (PGRP-LB, PGRP-SC2, PGRP-LF, and Pirk) of IMD across experimental groups, a noticeable trend emerges where *D. melanogaster* females exhibit higher upregulation compared to other groups; this might indicate that *D. melanogaster* females activated the IMD pathways at a level where negative regulation was initiated, whereas other experimental groups were unable to reach that elevated level. The reason behind the significant upregulation of PGRP-LF and Pirk in *D. simulans* remains unclear. This is particularly ambiguous considering the survival data, where it is evident that *D. simulans* was not effectively handling the infection at 16hpi (Figure 2-3, Figure 10-11, File S11).

Despite the upregulation of negative regulators, *D. melanogaster* females upregulate effector genes of the IMD pathway to a higher degree than males or *D. simulans;* this may reflect greater control of expression of effectors, as well as specific dynamics in female *D. melanogaster* in which upregulation of negative regulators is a response to the high levels of effector expression at 16hpi. This would then be expected to reduce effector expression at later time points, which was not possible to test from this experimental data. The key transcription factor Relish/NF-κB is upregulated less than males at 16hpi for *D. melanogaster* females; this might suggest there would be downregulation of the AMPs at later time points than those measured (Zaidman-Rémy et al. 2006). Eventually, the increased expression of genes involved in negative feedback loops regulating IMD in *D. melanogaster* females would be expected to result in reduced signaling and potentially reduced expression of other genes in the IMD pathway (Tanji and Ip 2005; Zaidman-Rémy et al. 2006; Vincent and Dionne 2021). These findings provide valuable insights into the intricate mechanisms underlying sex-specific immune responses and suggest a balance between gene activation and suppression as part of an adaptive strategy that contributes to increased survival in females.

### 4.4 Sex and species differences in the Toll pathway

Toll is another conserved signaling pathway and is involved in many functionally different processes, including innate immunity. Broadly, Toll-regulated AMPs were also upregulated in a female-biased manner in both species. The comparison between species revealed higher upregulation in *D. melanogaster* relative to *D. simulans,* which is also observed for IMD-regulated AMPs. However, the level of Toll-AMP upregulation is lower than it is for IMD-regulated AMPs in both sexes for *D. melanogaster* and *D. simulans.* This is expected because Toll is a more specialized response against gram-positive bacterial and fungal infections (Valanne et al. 2011). In the IMD pathway, the core transcription factors Relish showed sex differential expression for both species, but neither of the key transcription factors (Dif and Dorsal) in the Toll pathway were expressed differentially in either species. This suggests that the expression of Dif and Dorsal are not directly responsible for sex differences in the expression of AMPs, although sex differences may result from post-transcriptional regulation or differences in transcript levels prior to 16hpi (Schlamp et al., 2021). In addition, the Toll-associated AMPs might be regulated by other noncanonical transcriptional mechanisms to elicit sex dimorphism (Tanji and Ip 2005).

Sex differential regulation was detected for signal receptors and cellular response genes for both species. Among these genes involved in *spz* activation, PGRP-SA, PGRP-SD, and Spirit were upregulated in females and males of both species. This is expected to be associated with PRR-dependent Toll activation. In addition, Clip-SP Hayan, one of the serine proteases of the melanization cascade, was significantly overexpressed in females of both species relative to males (*p*-adj < 0.001) (Figure S5, File S11). The analysis also detected a notable trend where females (7-Fold) and males (4-Fold) of *D. simulans* drive significant upregulation of Clip-SP Psh (File S10). Both Clip-SP Hayan and Clip-SP Psh play a vital role in regulating the Toll pathway downstream of Clip-SP Grass, ModSP, and PRRs (Yamamoto-Hino and Goto 2016; Issa et al. 2018; Dudzic et al. 2019).

Persephone, a circulating serine protease, functions as an immune sensor for microbial proteolytic activities upstream of the Toll pathway by detecting exogenous proteases (Chamy et al. 2008; Issa et al. 2018). Upon detection, Persephone can elicit Toll activation in two different ways, through Psh-SPE-Spz or Psh-Protease-Spz (SPE independent) activation (Ligoxygakis et al. 2002; Gottar et al. 2006; Chamy et al. 2008; Yamamoto-Hino and Goto 2016). Although it is not established that Hayan can detect microbial proteases, it plays a vital role in both PRR-independent activation mechanisms (Dudzic et al. 2019). Previous studies showed that animals with SPE null mutations can perform the pro-Spz to Spz cleavage, suggesting that another SP can cleave Spz beyond SPE (Yamamoto-Hino and Goto 2016; Dudzic et al. 2019).

It is also possible that Hayan is upregulated to promote the melanization cascade, but the other key members of the Hayan-dependent blackening reaction, PPO1, and PPO2, were downregulated in *D. simulans* (Binggeli et al. 2014; Dudzic et al. 2015, 2019). In addition, the regulation of Hayan is strictly controlled by the serpins (Spn) for melanization (Binggeli et al. 2014). Analysis of the negative regulators of the melanization cascade reveals that Spn28Dc, a member of the Serpin family, which can negatively regulate the melanization process, is significantly upregulated in females and males of *D. simulans* and *D. melanogaster* (Ligoxygakis 2002; Tang 2009; Binggeli et al. 2014). This indicates that Hayan is more likely to be involved in the PRR-independent Toll activation rather than melanization.

Based on the observation, we hypothesize that significant downregulation of SPE in females (log2FC = - 1.31) and males (log2FC = −0.44) in *D. simulans* coincides with the use of the alternate Toll activation pathways, whereas females (log2FC = 0.91) and males (log2FC = 0.65) of *D. melanogaster* might sufficiently upregulate SPE; thus *D. melanogaster* is not depending on the microbial proteases-based Psh-SPE-Spz or Psh-Protease-Spz (SPE independent) activation at 16hpi (Issa et al. 2018; Dudzic et al. 2019). In contrast, Toll responses in *D. simulans* might be enhanced by the additional microbial protease based Psh-SPE-Spz or Psh-Protease-Spz (SPE independent) activation allowing maintenance of appropriate Spz levels (Yamamoto-Hino and Goto 2016; Issa et al. 2018; Dudzic et al. 2019). We note that it is not possible to make conclusions from our single timepoint data, although our results support the investigation of the hypothesis that SPE at 16hpi is downregulated, with *D. simulans* flies using Psh-Protease-Spz (SPE independent) activation.

Immune responses incur significant energetic costs for organisms. Therefore, the success of an organism depends not only on the activation of immune pathways but also on their effective suppression. Among the suppressors of Spz activation and the Toll pathway, two of the Serpins family members named Spn88Ea and Spn42Dd were overexpressed in both sexes of *D. melanogaster* (*p*-adj < 0.001). The role of the Spn88Ea and Spn42Dd is to negatively regulate the production of antifungal peptides via the Toll pathway (Reichhart et al. 2011). As mentioned earlier, there is less expression of the fungal-related AMPs except for *Drosomycin (Drs),* which has multiple roles as antipathogenic peptides, including function against bacterial infection (Hanson and Lemaitre 2020; Lin et al. 2020). These findings also explain the downregulation of the fungal cell wall pattern recognition receptor GNBP3 in the infected females (log2FC = −0.60) and males (−0.61) relative to wound control in *D. melanogaster* (Figure S2, File S10).

## 5. Conclusion

Our data showed that similar genes and pathways are upregulated in males and females, but the degree of upregulation differs between sexes and species. The comparative analysis revealed that these sex differences in the upregulation of immune-related biological processes are conserved between the two closely related species examined; interestingly, the sexes appear to suppress different sets of genes, indicating potential differences in trade-offs and resulting strategies. However, there are differences between species, both *D. melanogaster* sexes show higher upregulation of immune genes relative to *D. simulans*. Moreover, among the four experimental groups, the *D. melanogaster* female upregulated immune genes at the highest level relative to males and *D. simulans* of both sexes – which is concordant with the differences observed in survival. The downregulation of biological processes in females and males showed significant differences in both species. Although the pattern of downregulated GO terms is similar in males between species, the categories corresponding to downregulated genes in females are significantly different.

It has been reported in prior studies that Toll and IMD are responsible for the sex differences in *D. melanogaster*, and our results showed there were sex differences in the regulation of both conserved bacterial infection response pathways, IMD and Toll, which supports these previous studies (Duneau et al. 2017b; Vincent and Dionne 2021). Intriguingly, our data strongly suggest that *D. melanogaster* and *D. simulans* employ two mechanisms to activate Toll pathways differently during infection by *P. rettgeri*, with *D. melanogaster* using the canonical PRR-based Toll activation, and *D. simulans,* the microbial protease recognition-based Persephone pathway to stimulate Toll effectors along with the canonical path. The AMP expression level of *D. melanogaster* is consistent with the hypothesis that PRR-SPE-Spz activation alone can mount a heightened immune response in *D. melanogaster* relative to both processes in *D. simulans*. The IMD pathway genes showed sex-differential expression and varied in the degree of expression across species. The analysis of the negative regulators showed *D. melanogaster* females were upregulating four key “brakes” (PGRP-LB, PGRP-SC2, PGRP-LF, and Pirk) to prevent the overstimulation of IMD pathways at 16hpi. Overall, our finding suggests that sex differences in the expression of immune genes are typically conserved in the two species, for most of the upregulated genes with the key exception of negative regulators but noticeably different for downregulation; also, there were significant sex differences in the fold-change of up-or downregulation between *D. melanogaster* and *D. simulans*.

This study will significantly contribute to our understanding of the evolution of sex differences and point to specific testable hypotheses that might explain observed species and sex differences in survival. Further research involving both within and between-species comparisons is essential to ascertain whether these differences are consistent across various contexts. The observed variation in infection outcomes among different reports, even when using the same species and pathogens, emphasizes the significance of considering multiple sources of variation within individual studies. Additionally, conducting comparisons across a diverse range of time points, rather than relying on a single snapshot, would provide a more comprehensive view of sex dimorphisms occurring at different stages of disease progression.

## Data availability

All raw and processed sequencing data generated in this study can be found on NCBI Gene Expression Omnibus (GEO; https://www.ncbi.nlm.nih.gov/geo/) (in processing). The authors confirm that all data supporting this article’s conclusions are included in the article, figures, supplement figures, and supplement tables.

## Acknowledgments

We extend our gratitude to the Bloomington Stock Center for providing the flies. We also thank Brian Lazzaro from the Department of Ecology and Evolutionary Biology at Cornell University for supplying the bacterial strain. Our appreciation goes to Blake Rochester, Bayley Atkins, Isabella Blair, and Audrey Powell for their assistance with fly work, and to the other members of the Graze lab for their valuable discussions. RNA sequencing was conducted at the Genomic Sequencing and Analysis Facility, University of Texas at Austin.

---

**Figure S1:**
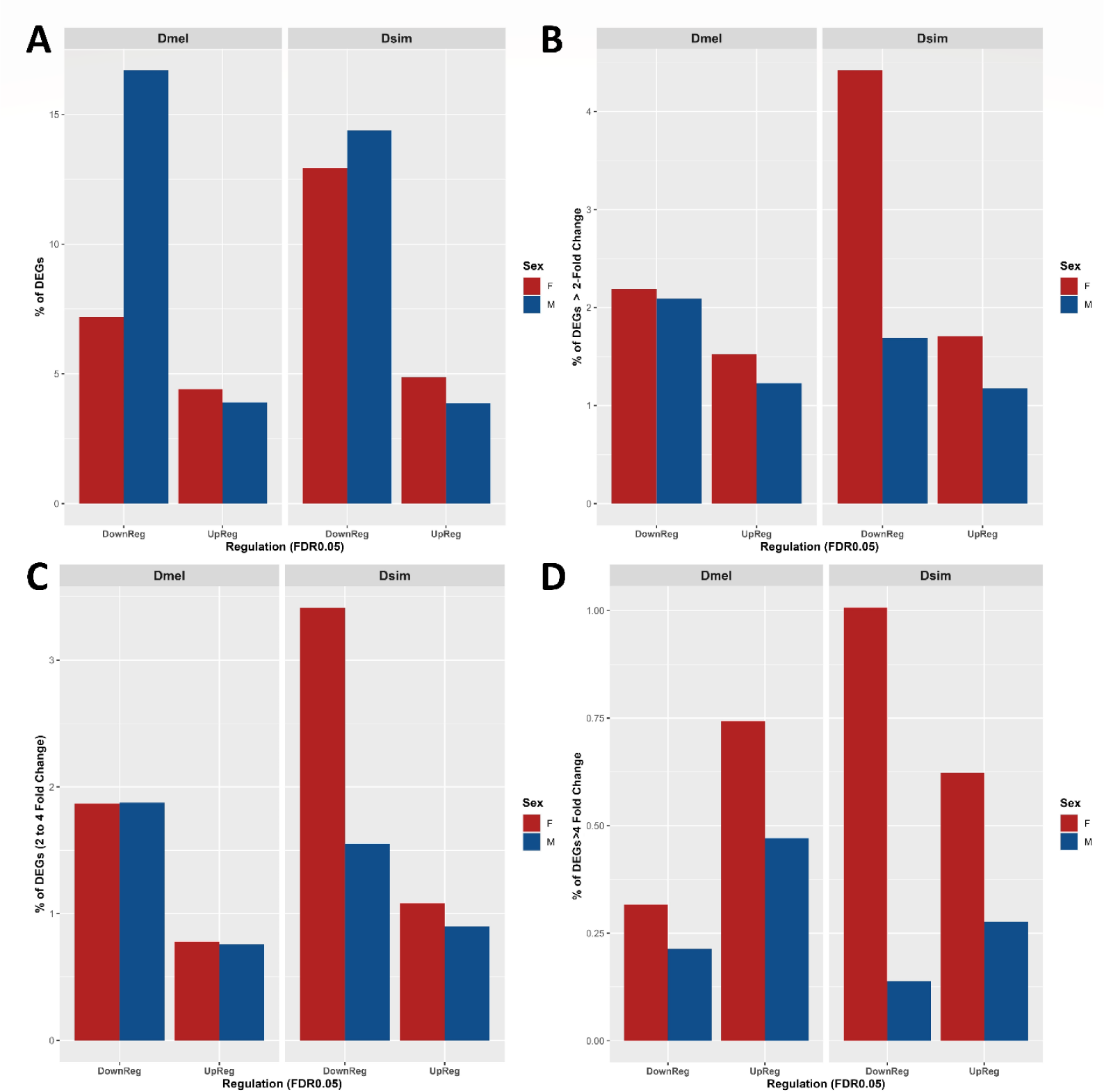
Percent of differentially expressed gene (DEGs) in the females (F) and males (M) in *D. melanogaster* (Dmel) and *D. simulans* (Dsim) upon *P. rettgeri* (PRET) infection for different ranges of fold change (FC). **A.** Percentage of up- and downregulated DEGs (without FC cutoff), **B.** Percentage of up- and downregulated DEGs at > 2FC**, C.** Percentage of up- and downregulated DEGs at 2 to 4FC, **D.** Percentage of up and downregulated DEGs at > 4FC. The percentage of expressed genes showed differential expression with cut-off FDR < 0.05 when compared to the wound control (PBS). The expressed genes were defined after filtering for low expression, the number of expressed genes was 13588 and 13010 for *D. melanogaster* and *D. simulans*, respectively (for details, File S4-S5). The percent count of the more than 2-FC downregulated genes showed the difference between females and males in *D. melanogaster* is much more comparable, in contrast to the pattern for downregulation overall (without FC) (Figure S1). In *D. simulans,* downregulation at more than 2-FC showed the opposite direction than overall downregulation (without FC). For the greater than 2-FC cutoff, upregulation is comparable in both species for females and males. For genes with the largest differences in expression, > 4 FC, in *D. simulans*, many more genes are downregulated in females than in males, and a similar pattern is revealed in upregulation. These suggest that while overall downregulation was more prevalent in males for *D. simulans*, when the degree of suppression is dependent on sex, females showed more highly downregulated genes (at > 2-FC, 2-FC < FC < 4-FC, > 4-FC). This is also true for the *D. melanogaster* (at > 2-FC and > 4-FC), although the magnitude of the difference is not as high as in *D. simulans* (Figure S1).

**Figure S2:**
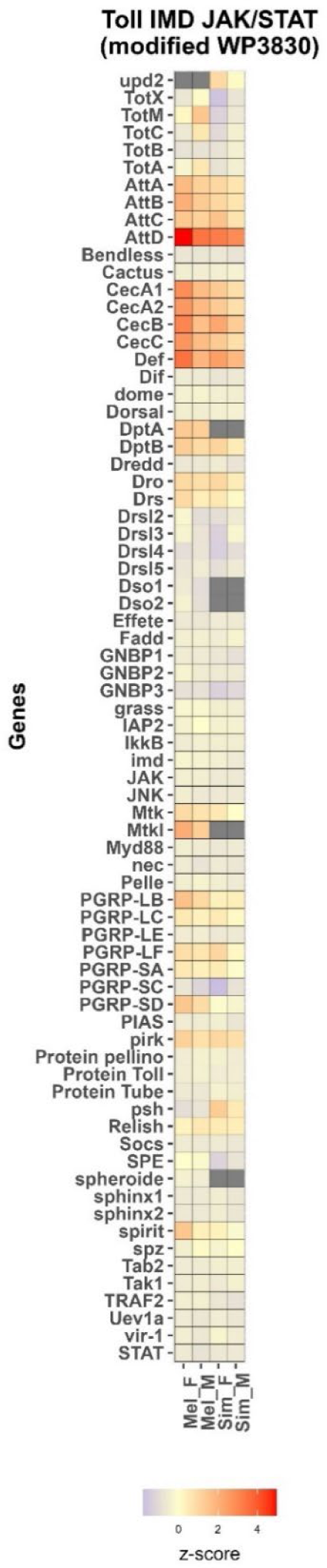
Sex differences in the Toll-IMD-Jak/Stat signaling pathway (wiki pathway ID: WP3830) in *D. melanogaster* and *D. simulans* upon *P. rettgeri* infection. Mel=*D. melanogaster*, Sim=*D. simulans*, F=Female, M= Male. The expression (z-score) of the pathway was represented without statistical cut-off (for details, File S10).

**Figure S3:**
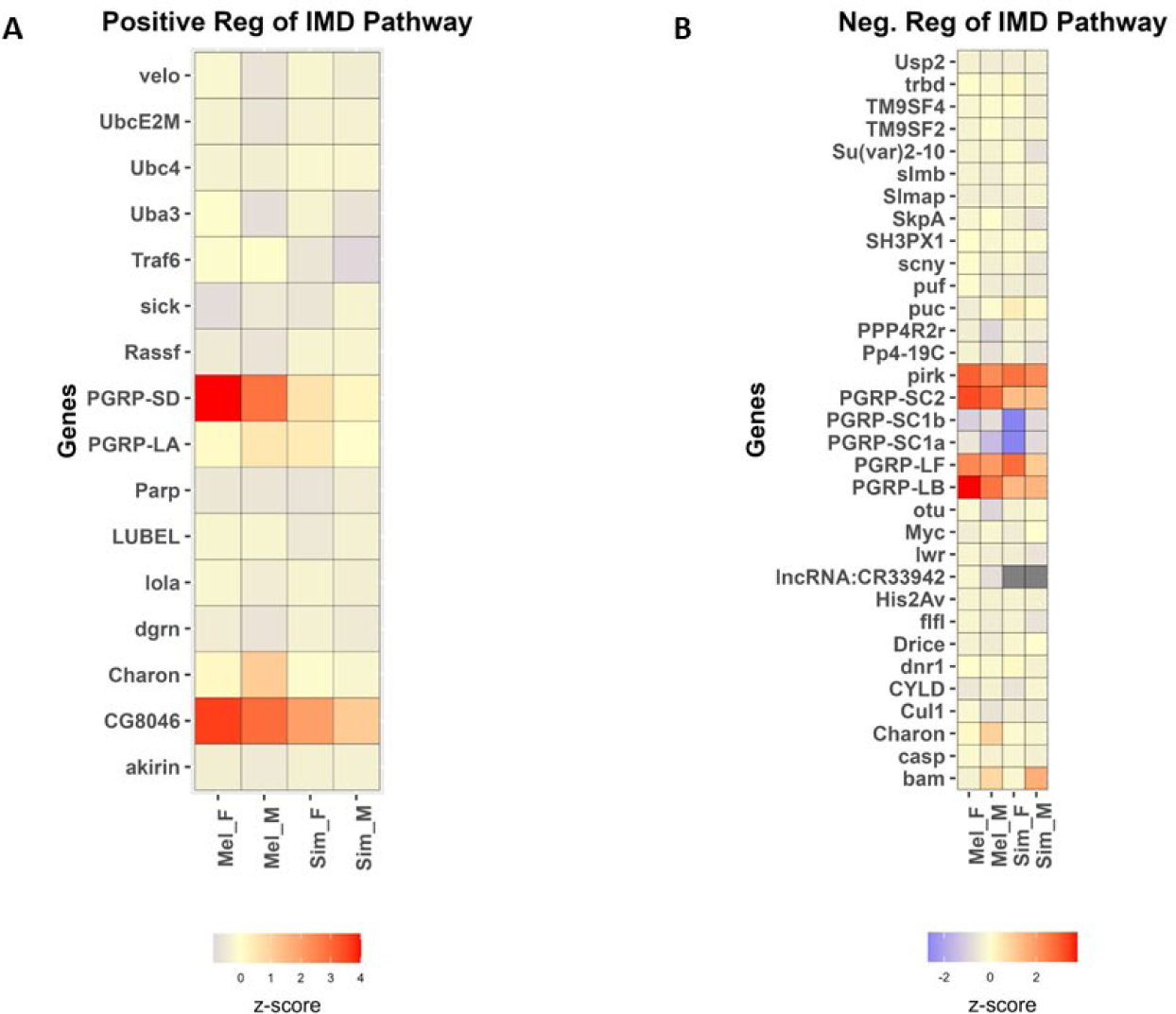
Sex differences in **A.** positive and **B.** negative regulators of the IMD pathways in *D. melanogaster* and *D. simulans* upon *P. rettgeri* infection. Mel=*D. melanogaster*, Sim=*D. simulans*, F=Female, M= Male. The expression (z-score) of the pathway was represented without statistical cut-off (for details, File S11).

**Figure S4:**
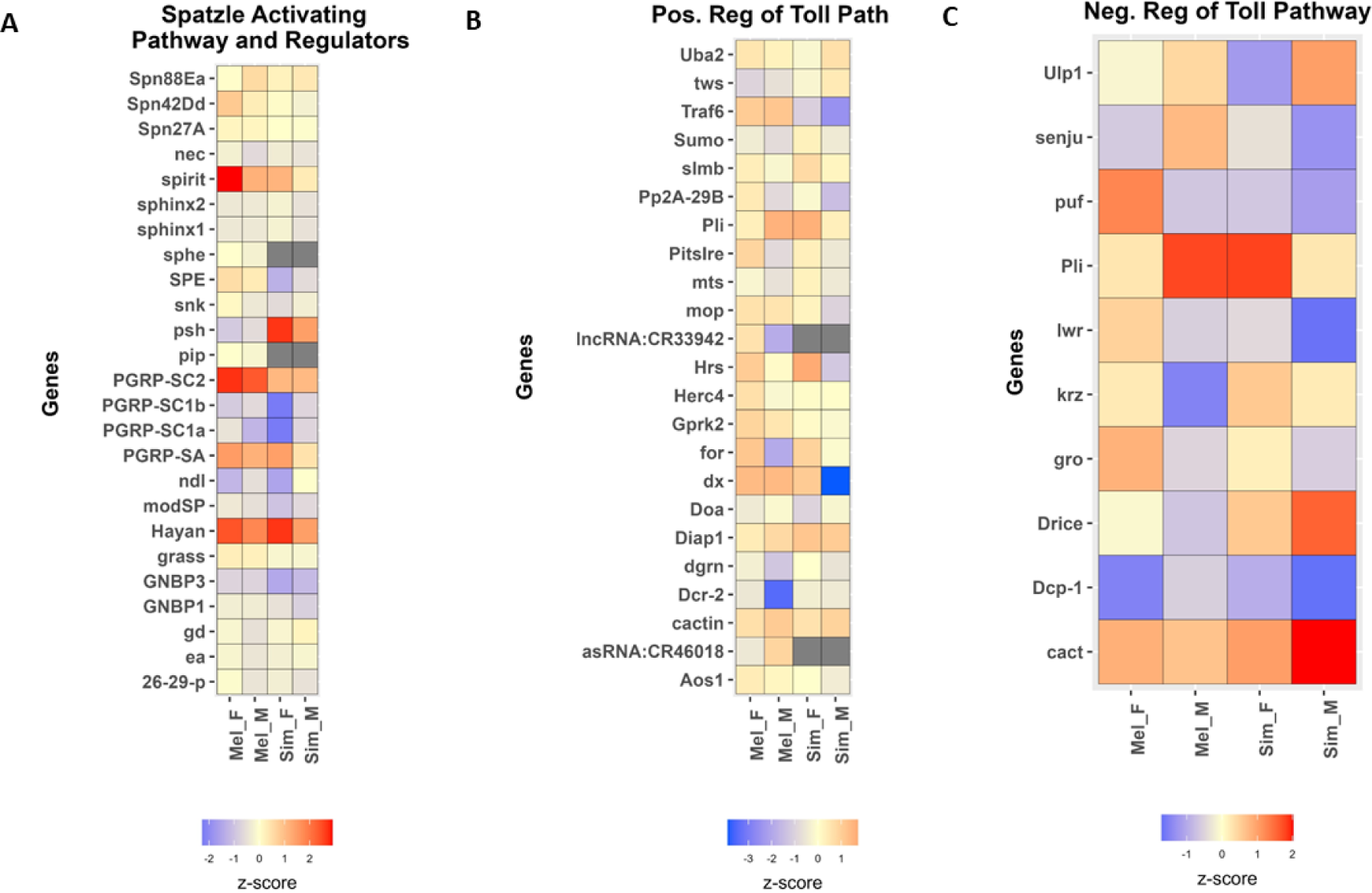
Sex differences in **A.** *spz* activating pathway, **B.** positive and **C.** negative regulators of the Toll pathways in *D. melanogaster* and *D. simulans* upon *P. rettgeri* infection. Mel=*D. melanogaster*, Sim=*D. simulans*, F=Female, M= Male. The expression (z-score) of the pathway was represented without statistical cut-off (for details, File S11).

**Figure S5:**
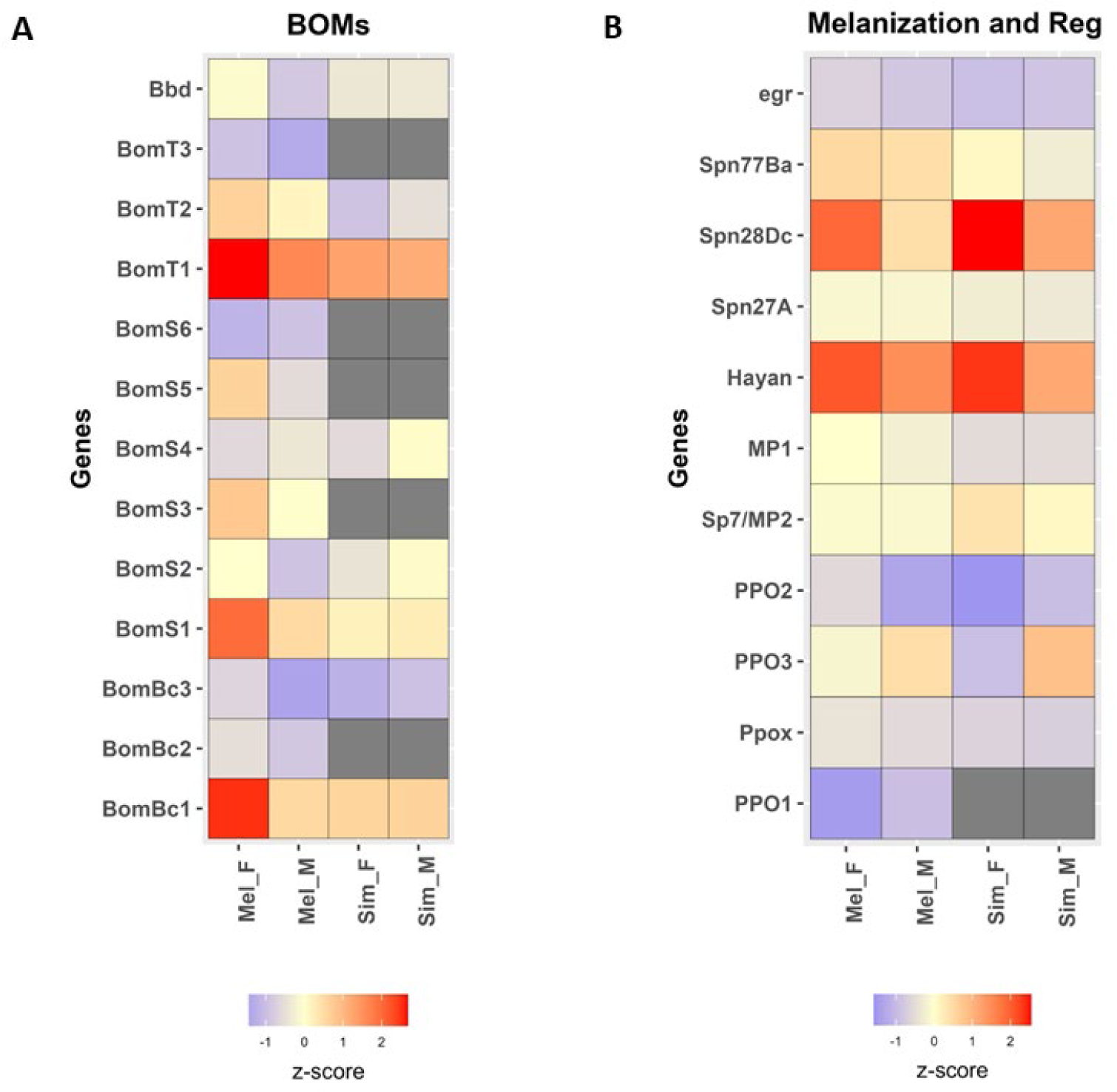
Sex differences in **A.** Bomanins (Boms) and **B.** melanization in *D. melanogaster* and *D. simulans* upon *P. rettgeri* infection. Mel=*D. melanogaster*, Sim=*D. simulans*, F=Female, M= Male. The expression (z-score) of the pathway is represented without statistical cut-off (for details, File S11).

**Figure S6:**
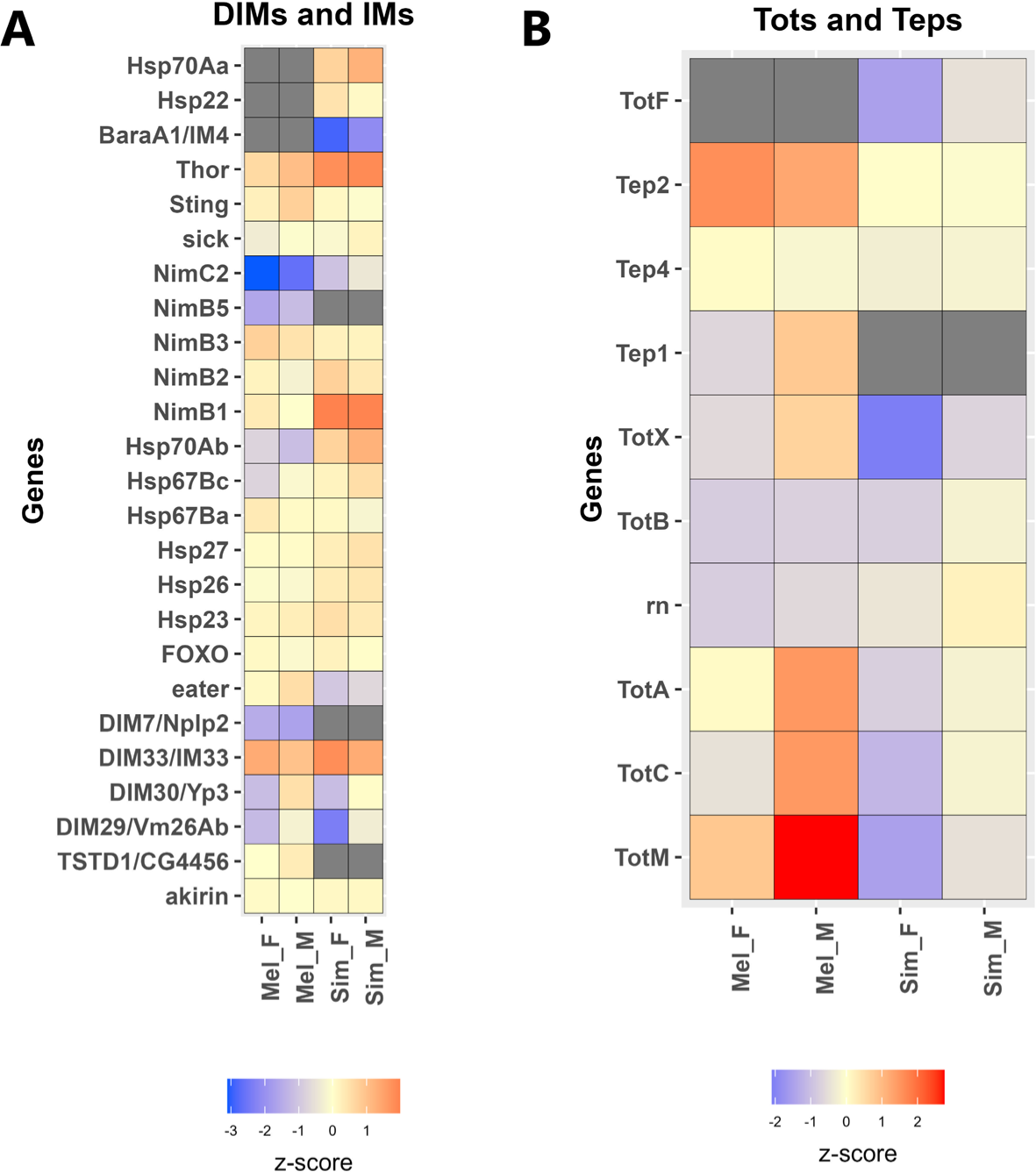
Sex differences in **A.** DIMS and IMs, and **B.** Tots, and Teps in *D. melanogaster* and *D. simulans* upon *P. rettgeri* infection. Mel= *D. melanogaster*, Sim= *D. simulans*, F= Female, M= Male, DIMs= Drosophila immune-induced molecules, IMs= Immune Molecules, Tots= Turandot, Teps= Thioester-containing protein, AMPs= Antimicrobial peptides. The expression (z-score) of the pathway was represented without statistical cut-off (for details, File S11).

**Figure S7:**
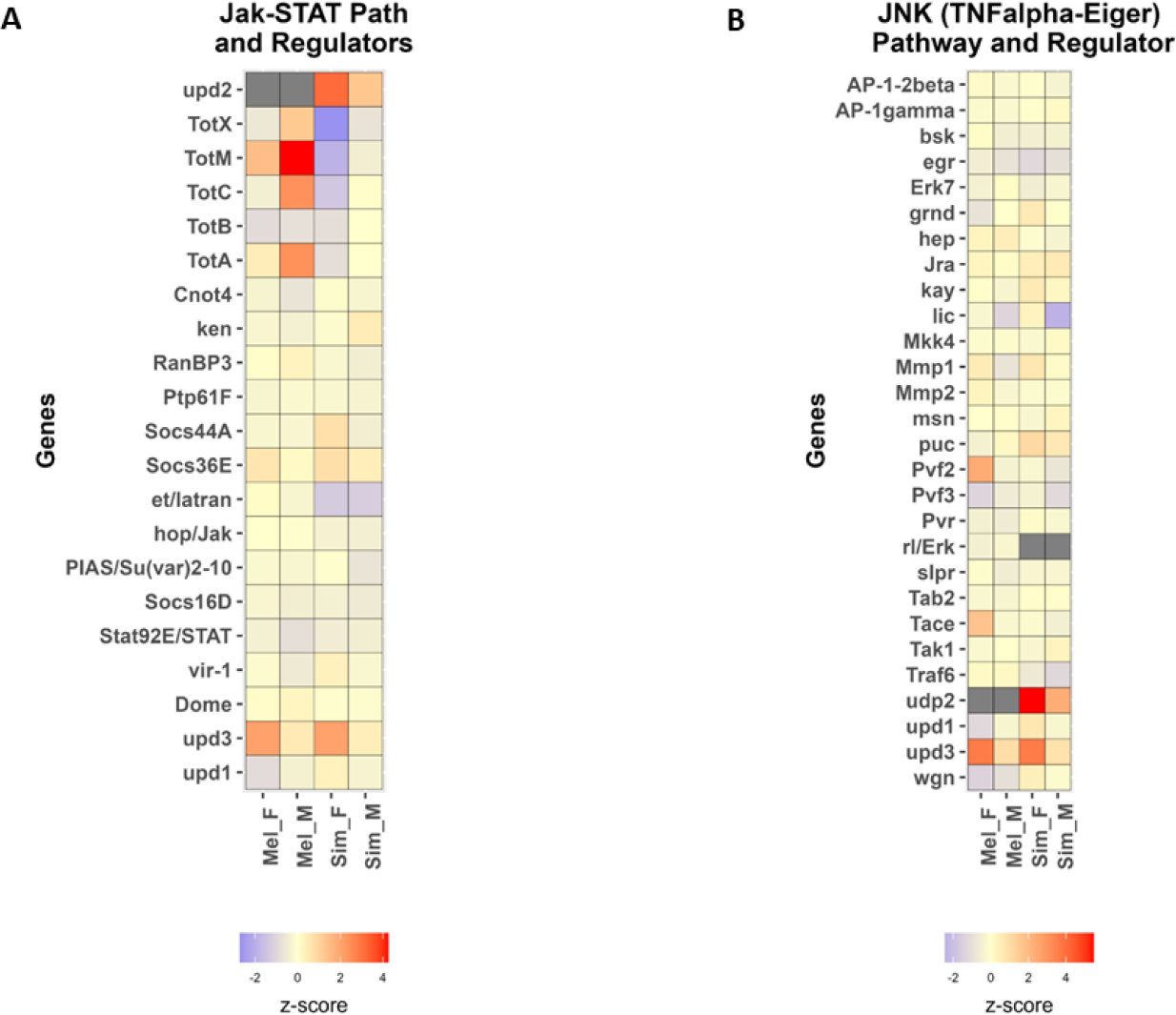
Sex differences in **A.** JAK-STAT (hop-STAT92E) and **B.** Jnk (TNFa-Eiger) pathway with regulators in *D. melanogaster* and *D. simulans* upon *P. rettgeri* infection. Mel=*D. melanogaster*, Sim=*D. simulans*, F=Female, M= Male. The expression (z-score) of the pathway was represented without statistical cut-off (for detail, File S10-S11).

**Table S1:**
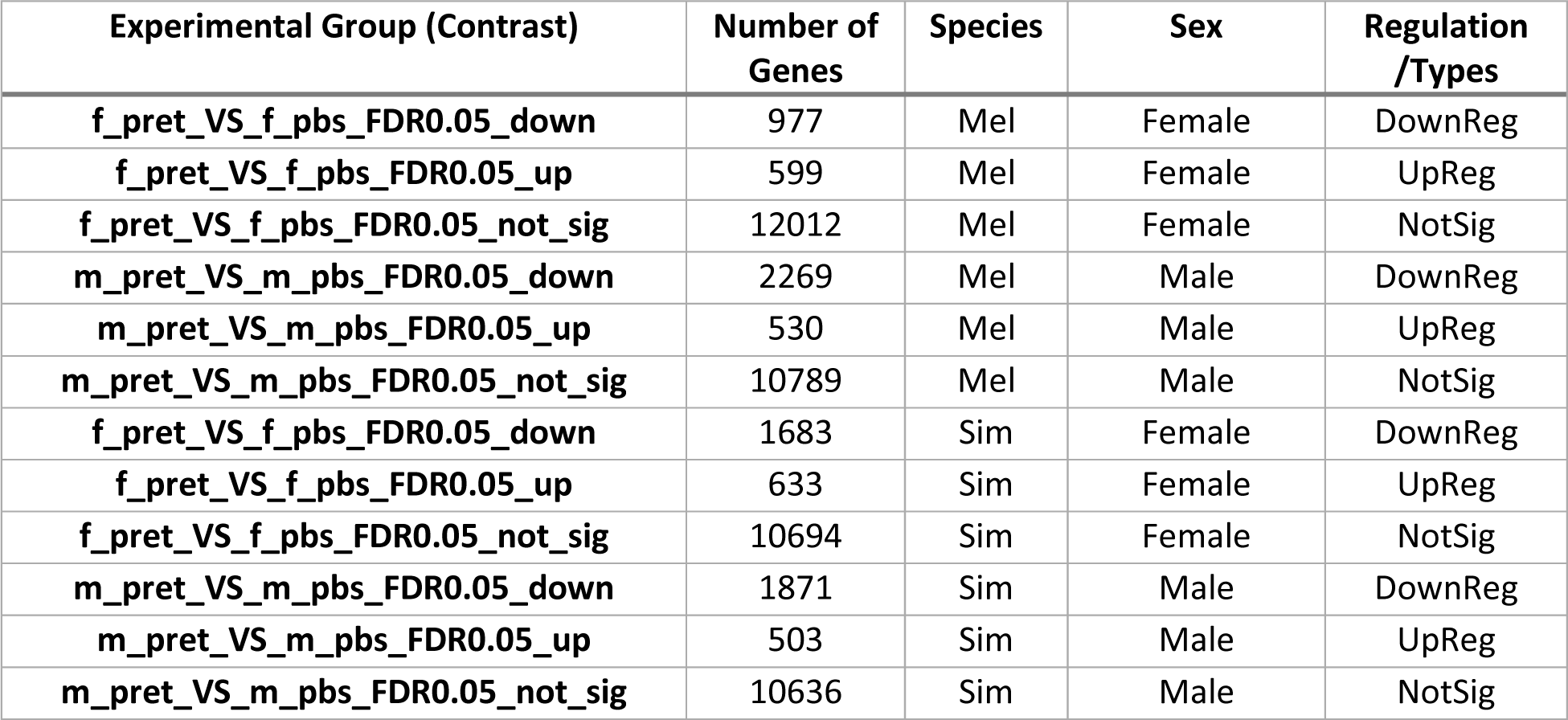
The number of genes upregulated, downregulated, and not significant at FDR < 0.05 for *D. melanogaster* and *D. simulans*. f=Female, m=make, pbs=phosphate buffer solution, pret= *P. rettgeri*, Mel= *D. melanogaster*, Sim= *D. simulans*, DownReg= Down-Regulation, UpReg= Up-Regulation, NotSig= Not significant, FDR= False Discovery Rate (for details, File S4-S5).

**Table S2:** Up- and down-regulated enriched gene ontology terms biological process (GO: BP) in males and females. Mel=*D. melanogaster*, Sim=*D. simulans*, Com. En. w= Commonly Enriched with, UpReg=Up Regulation, DwReg=Down Regulation (for details, File S9).

| Exp Groups | Total Enriched | Up Reg | Com. En. w Opposite Sex-UpReg | Com. En. w compared Species-UpReg | Dn Reg | Com. En. w Opposite Sex-DnReg | Com. En. w compared Species-DnReg |
| --- | --- | --- | --- | --- | --- | --- | --- |
| Mel Female | 50 | 49 | 46 | 44 | 1 | 1 | 0 |
| Mel Male | 101 | 84 | 46 | 56 | 17 | 1 | 12 |
| Sim Female | 68 | 47 | 42 | 44 | 21 | 0 | 0 |
| Sim Male | 72 | 57 | 42 | 56 | 15 | 0 | 12 |

